# FOXA1/2 depletion drives global reprogramming of differentiation state and metabolism in a human liver cell line and inhibits differentiation of human stem cell-derived hepatic progenitor cells

**DOI:** 10.1101/2020.06.01.128108

**Authors:** Iyan Warren, Michael M. Moeller, Daniel Guiggey, Alexander Chiang, Mitchell Maloy, Ogechi Ogoke, Theodore Groth, Tala Mon, Saber Meamardoost, Xiaojun Liu, Sarah Thompson, Antoni Szeglowski, Ryan Thompson, Peter Chen, Ramasamy Paulmurugan, Martin L. Yarmush, Srivatsan Kidambi, Natesh Parashurama

## Abstract

FOXA factors are critical members of the developmental gene regulatory network (GRN) composed of master transcription factors (TF) which regulate murine cell fate and metabolism in the gut and liver. How FOXA dictates human liver cell fate, differentiation, and simultaneously regulate metabolic pathways is poorly understood. Here, we aimed to determine the role of FOXA2 (and FOXA1 which is believed to compensate for FOXA2) in hepatic differentiation and cell metabolism in a human hepatic cell line (HepG2). siRNA targeting of FOXA1 and FOXA2 in human hepatic (HepG2) cells and during hepatic differentiation significantly downregulated albumin (p < 0.05) and GRN TF gene expression (HNF4A, HEX, HNF1B, TBX3) (p < 0.05) and significantly upregulated endoderm/gut/hepatic endoderm markers (goosecoid (GSC), FOXA3, and GATA4), gut TF (CDX2), pluripotent TF (NANOG), and neuroectodermal TF (PAX6) (p < 0.05), all consistent with a partial/transient cell reprogramming. shFOXA1/2 targeting resulted in similar findings and demonstrated evidence of reversibility. RNA-seq followed by bioinformatic analysis of shFOXA1/2 knockdown HepG2 cells demonstrated 235 significant downregulated genes and 448 upregulated genes, including upregulation of markers for alternate germ layers lineages (cardiac, endothelial, muscle) and neurectoderm (eye, neural). We found widespread downregulation of glycolysis, citric acid cycle, mitochondrial genes, and alterations in lipid metabolism, pentose phosphate pathway, and ketogenesis. Functional metabolic analysis agreed with these findings, demonstrating significantly diminished glycolysis and mitochondrial respiration, and accumulation of lipid droplets. We hypothesized that FOXA1/2 inhibit the initiation of human liver differentiation *in vitro*. During hPSC-hepatic differentiation, siRNA knockdown demonstrated de-differentiation and unexpectedly, activation of pluripotency factors and neuroectoderm. shRNA knockdown demonstrated similar results and activation of SOX9 (hepatobiliary). These results demonstrate complex effects of FOXA1/2 on hepatic GRN effecting de-differentiation and metabolism with applications in studies of cancer, differentiation, and organogenesis.

## INTRODUCTION

Hepatic gene expression is precisely controlled by regulatory transcriptional factors (TFs) which form core gene regulatory networks (GRN). The well-studied hepatic GRN establish and maintain liver differentiated function, and include FOXA1, FOXA2, FOXA3, HNF1A, HNF1B, HNF4A, HNF6, HEX, TBX3, and PROX1 ^1–2^ . They mediate not only differentiation and lineage choice ^3^, but also key aspects of metabolism ^4^, and regeneration ^5^. Hepatic GRN also play a role in physiology and pathophysiology ^6–7^, and defining disease states like hepatic cancer ^8^, fibrosis ^9^, and alcoholic hepatitis ^10^. Finally, TF-based reprogramming experiments also deploy hepatic GRN in fibroblasts reprogrammed to hepatocyte-like cells ^11^. Thus, hepatic GRN have broad significance in controlling differentiation state, metabolism, and pathophysiological processes.

FOXA factors are members of the hepatic GRN and play a key role in controlling liver differentiation. Mouse genetic studies of FOXA2, and its compensating factor FOXA1, demonstrate stage-specific developmental roles including: 1) foregut endoderm induction ^12–13^ 2) definitive endoderm maintenance ^14^, 3) initiating liver development ^15^ and 4) maintaining hepatic gene expression in adult liver (FOXA1/2/3) ^16^. Seminal studies demonstrates FOXA2 functions as a “pioneer factor” that binds to and remodels compacted, normally inaccessible, chromatin at the silent ALB promoter/enhancers leading to assembly of liver TFs and ALB gene expression ^17–21^ . FOXA1 compensates for FOXA2, which requires a double knockout to prevent redundant function. The FOXA1/2 phenotype has been investigated in murine bile ducts, pancreas, the lung, and intestinal differentiation ^22–25^ . FOXA1/2 plays a role in initiating liver differentiation, but it has been found that for maintaining mature liver, FOXA3 is also required ^16^. FOXA3 maintains enhancer activity, chromatin accessibility, nucleosome positioning, and binding of HNF4a, and FOXA1/2/3 -/-/-phenotype results in hepatocyte de-differentiation towards endoderm ^16^. Taken together, FOXA1 compensates for FOXA2 during the initiation of murine liver development, and FOXA1/2/3 are all involved in maintaining developmentally regulated liver gene expression in murine adult liver, and FOXA3 compensates for the loss of FOXA1/2.

The liver GRN not only control differentiation, they also play a key role in normal metabolism and disease. They regulate metabolism within hepatocytes, including carbohydrate (glucose) metabolism, glutamine metabolism, nitrogen (urea) metabolism, lipid metabolism, glycogen storage and glycogenolysis, gluconeogenesis, amino acid metabolism, and control of fasted, fed, and inflammatory states ^26^. Early studies include showed FOXA2 regulates lipid, carbohydrate, xenobiotic metabolism and demonstrate global binding of liver-specific proteins ^20, 27^. As part of the hepatic GRN, FOXA factors often partner with HNF1A, and HNF4, which togethers play a role in congenial diabetes ^28–30^ and fatty liver disease ^31^. FOXA factors also play a key role hepatocellular carcinoma (HCC) and liver fibrosis. ^32–35^

Despite extensive murine studies, the role of FOXA in human hepatocyte metabolism and human liver cell lines remains far less understood ^1^. The regulation of human GRN by FOXA factors has been investigated using models of human liver cell lines and models of human pluripotent stem cell (hPSC)-liver differentiation. In CRISPRi screens during hPSC endoderm differentiation, FOXA2 controls differentiation ^36^. Further, FOXA1/2 primes human liver genes within hPSC-derived gut tube endoderm together with corresponding epigenetic changes in histone methylation ^37–38^. Further, FOXA1/2 has been shown to bind cooperatively with FOXO in regulating Insulin growth factor (IGF) binding protein 1 in human HEPG2 cells, demonstrating a link to metabolism ^39^. Nearly all liver cell line studies (HepG2 cells) ^40^ do not formally target both FOXA1/2. Further, more human studies are critical, since human GRN that govern development bind to genes different than from murine ones ^41^. Taken together, how FOXA initiates human hepatic differentiation is poorly understood. Further there remain gaps in our understanding of FOXA1/2 in regulating liver GRN, differentiation, and metabolism in human liver cell lines and human hPSC-derived gut tube endoderm progenitors.

We hypothesized that FOXA1/2 knockdown effects both differentiation state and metabolism. We found that that siRNA and shRNA-mediated knockdown of FOXA1/2 in a human liver cell line (HepG2) results significant downregulation of ALB expression. Further, we observed global downregulation of all major liver GRN, and unexpectedly, upregulation of alternate germ layer master factors, including pluripotency, neural, cardiac, and endoderm genes. Transcriptomic and bioinformatic analysis confirmed these changes in both differentiation and metabolism, and functional metabolic analysis confirmed extensive metabolic changes. We then tested the role FOXA1/2 in a hPSC-hepatic differentiation model. The data shows that FOXA1/2 controls the initiation of human liver differentiation, in part through liver GRN. Therefore, FOXA factors could be therapeutic targets for understanding fibrosis, and cancer, and may function as pharmaceutical targets to favorable control these processes *in vitro* or *in vivo*.

## METHODS

### Reagents/Materials

Dulbecco’s modified Eagle’s medium (DMEM) (Cat. #: 10566024), OptiMEM (Cat. #: 31985070), RPMI (Cat. #: 61870036), IMDM (Cat. #: 12440053), KO Serum (10828010), Penicillin-streptomycin (10000 U/mL) (Cat. #: 15140122), Fetal Bovine Serum (Cat. #: A3160602), 0.05% Trypsin-EDTA (Cat. #: 25300062), B27 (containing insulin) (50x) (Cat. #: 17504044), Ham’s F12 (Cat. #: 11765054), N2 Supplement (100x) (Cat. #:17502048), Lipofectamine RNAiMAX Transfection Reagent (Cat. #: 13778030), Beta Actin Loading Control Antibody (Cat. #: MA5-15739), Puromycin: Liquid (20mL) (Cat. #: A1113802), High glucose DMEM (Invitrogen Cat. #: 10566024), DAPI (Cat. #: 62248), Precast gels-Bolt 4-12% Bis-Tris Plus 10-well gels (Cat. #: NW04120BOX), and nitrocellulose transfer stacks (Cat. #: IB23002) were purchased from ThermoFisher (Waltham, MA). Activin A (Cat. #: 78001.1), mTESR1 (Cat. #: 85850), keratinocyte growth factor (KGF, FGF-7) (Cat. #: 78046), fibroblast growth factor 2 (FGF-2, bFGF) (Cat. #: 78003.1), epidermal growth factor (EGF) (Cat. #: 78006.1), and Gentle Cell Dissociation Reagent (GDR) (Cat: #: 07174), were purchased from StemCell Technologies (Vancouver, CA). G418 (Cat. #: G8168-10ML), Doxycycline, (Cat. #: D9891-1G), L-Ascorbic Acid (Cat. #: A4544), MTG (Monothioglycerol) (Cat. #: M6145), Polybrene (hexadimethrine bromine) (Cat. #: 107689), were purchased from Sigma Aldrich (St. Louis, MO). Aurum Total RNA Mini Kit (Cat. #: 7326820), DNase I (Cat. #: 7326828), iTaq Universal SYBR Green Supermix (Cat. #: 1725121), and iScript cDNA Synthesis Kit (Cat. #: 1708891) were purchased from Bio-Rad. Pooled siRNA for FOXA1 (siGENOME SMART POOL Human FOXA2 (#3170)) and for FOXA2 (siGENOME SMARTPOOL Human FOXA1 (#3169)) were purchased from Dharmacon (now Horizon Discovery Group (Waterbeach, UK). siGLO Cyclophilin B Control siRNA (D-001610-01-05) and non-targeting siGENOME nontargeting pool #1 (D-0012060-13-05) was purchased from Dharmacon GE life sciences (now Horizon Discovery Group (Waterbeach, UK). We also used Foxa1 silencer select siRNA, Cat #: 4392420, Assay ID: s6689, Foxa2 silencer select siRNA, Cat #: 4392420, Assay ID: s6692, Silencer select siRNA GAPDH positive control, Cat #: 4390849, Silencer select siRNA negative control #1, Cat #: 4390843 from Thermofisher. Matrigel, Growth factor-free (Cat. #: 40230), was purchased from Corning (Corning, NY). Bovine Serum Albumin (Cat. #: 10791-790) was purchased from VWR International (Radnor, PA). CHIR (CHIR99021) (Cat. #: 13122) was purchased from Cayman Chemical (Ann Arbor, MI). Rho-associated kinase (ROCK) inhibitor (Y27632 2, Cat. #: MBS577605) was purchased from Mybiosource.com. Fugene HD Transfection Reagent (Cat. #: E2311) was purchased from Promega (Madison, WI). 96-well PCR plates (Cat. #: L223080), tissue culture treated 24-well plate (Cat. #: 702001), 75-cm^2^ polystyrene tissue culture-treated flasks (Cat. #: 708003), PCR Plate Covers (Cat. #: HOTS-100) were purchased from Laboratory Product Sales, Inc (Rochester, NY). The pLKO.1 puro (Cat. #: 8453), pLKO-shFOXA1#1 (Cat. #: 70095), pLKO-shFOXA1#2 (Cat. #: 70096), and pLKO-shScramble. (Cat. #: 1864) plasmids were purchased from Addgene (Watertown, MA). Human hepatoma (HepG2) (Cat. #: HB-8065) were purchased from ATCC (Manassas, VA). All primers were purchased from either Integrated DNA technologies (IDT) (Newark, NJ), Sigma Aldrich (St. Louis, MO) or Thermofisher (Waltham, MA).

### Antibodies

Mouse anti-human beta actin (Cat. #: MA5-15739), mouse anti-human FOXA2 monoclonal (Cat. #: MA5-15542), Goat anti-mouse secondary antibody (Alexa Fluor 488) (Cat. #: A-11034), and Goat anti-rabbit secondary antibody (Alexa Fluor 568) (Cat. #: A-11011) were purchased from ThermoFisher (Waltham, MA). Mouse anti-human ALB monoclonal (Cat. #: ab10241) was purchased from Abcam (Cambridge, MA). Mouse anti-human albumin monoclonal antibody (Cat. #: sc-271604), mouse anti-human AFP monoclonal antibody (Cat. #: sc-130302), Mouse anti-human OCT3/4 antibody (Cat. #: sc-5279), FITC- and TRITC-conjugated, anti-mouse, rabbit, and goat antibodies, and IgG control antibodies were purchased from Santa Cruz Biotechnology (Dallas, TX). Rabbit anti-human SOX17 antibody (Cat. #: NBP2-24568) was purchased from Novus Biologicals (St. Louis, MO). HRP-conjugated secondary antibodies were purchased from Jackson ImmunoResearch Laboratories (West Grove, PA).

### Liver cell line culture

Human hepatoma (HepG2) cells, derived from 14 y/o patient with hepatoblastoma (ATCC, Cat. #: HB-8065) and maintained with 10 mL of high glucose DMEM containing 1% Penicillin-Streptomycin (P/S), and 10% fetal bovine serum (FBS) in culture in 75 cm^2^ polystyrene tissue culture-treated flasks (LPS), and incubated at 5% CO_2_, 21% O_2_, and at 37°C. Medium changes were performed every two days. HepG2 cells were passaged weekly by the addition of 5 ml of either 0.05% or 0.25% Trypsin-EDTA, and replated at 1:5-1:10 dilution, with passage numbers for all experiments ranging between 15-35.

### Preparation of human pluripotent stem cell (hPSC) differentiation medium (SFD medium)

Basal differentiation medium for hPSC differentiation towards endoderm, gut tube, and liver contained SFD, a serum-free, defined medium based upon other mouse and human stem cell studies (Gadue, Huber et al. 2006). SFD medium contains 75% IMDM or RPMI supplemented with 25% Ham’s F12, 0.5% N2 Supplement, 0.5% B27 supplement, 2.5 ng/ml FGF2, 1% Penicillin + Streptomycin, 0.05% Bovine Serum Albumin, 2mM Glutamine, 0.5mM Ascorbic Acid, and 0.4mM Monothioglycerol (MTG).

### Feeder-free culture (maintenance) of human pluripotent stem cells (hPSCs)

We performed experiments using the UCSF4 human embryonic stem cell (hESC) line (NIH Registry 0044, female), a kind gift from Susan Fisher, PhD, Head of the Embryonic Stem Cell Center, UCSF Institute for Regenerative Medicine and stem cells. UCSF4 cells were karyotyped prior to use, and passage numbers used were 15-35. Further, a karyotyped, commercially available induced pluripotent stem cell (iPSC) line (BXS0114 ACS1028 female (ATCC) was expanded and used from passages 15-45. Human pluripotent stem cells (hPSC) were cultivated at 90% N_2_, 5% O_2_, 5% CO_2_, (Tri-gas HERAcell VIOS 160i CO_2_ incubators) using mTESR1 medium (warmed to room temperature), on 6-well tissue culture-treated plates, coated with 1:15 diluted (in DMEM) growth factor free-matrigel. Wells were coated with matrigel by adding 1 mL of diluted matrigel per well of a 6-well plate and incubating for 1.5-2 hrs. at 37°C. Excess dilute matrigel was then removed and the wells were washed with PBS. Cell culture medium was changed every other day. For passaging, the mTESR1 serum-free maintenance medium was removed, 1 mL of gentle cell dissociation reagent (GDR) (Stem Cell Technologies) was added for 10-15 minutes, and single cells or clumps of cells were harvested from the dish. Cells were centrifuged 3 minutes at 800-1000 RPM (Eppendorf 5810 tabletop centrifuge) and resuspended. Cells were frozen in mTESR1 medium + 5% DMSO at -80° C overnight, followed by liquid nitrogen cryostorage. Passage number varied between 15-35 for all experiments.

### Definitive endoderm induction from hPSC

hPSCs were harvested by replating cells in mTESR with 10 µM ROCK inhibitor overnight at 50,000-200,000 cells in a growth factor-free matrigel-coated, tissue culture treated 24-well plate. Coating was accomplished by adding 300 µL of diluted matrigel (1:15 growth factor-free matrigel in DMEM), per well, for 1.5-2 hours of coating, with excess matrigel removed. The next day, definitive endoderm was induced. In RPMI medium with 1x B27 (no insulin) and 0.2% KO serum, definitive endoderm was induced with Activin A (100 ng/ml) and CHIR (3 µM) for 1 day, followed by Activin A (100 ng/ml) for 3 more days. Medium was changed daily and 500-750 µL medium was used per well. The 0.2% KO serum was added for improved viability at 5% O_2_, and higher seeding densities improved culture morphology. Gut tube (GT) endoderm induction from definitive endoderm was performed in SFD medium (Gadue, Huber et al. 2006) (see: Preparation of SFD medium) containing keratinocyte growth factor (KGF or FGF7) (25 ng/ml) for an additional 2 days.

### Hepatic differentiation from human pluripotent stem cells (hPSC)

Two protocols were used for liver differentiation. *Growth factor-free gut tube/ hepatic differentiation protocol:* To promote spontaneous differentiation of hPSC under 5% O_2_ conditions, we added SFD medium with KGF (25 ng/ml) for a total of 10 days until day 14, with medium changes every day. Cells remained in 24-well plates, and SFD medium was changed daily. No additional growth factors were added. This is consistent with literature that in which mesoderm progenitors can stimulate liver differentiation in the absence of overt liver growth factors ^42–43^ and spontaneous hepatic differentiation of mouse ESC ^44^.

*Growth factor-positive hepatic differentiation protocol:* To promote hepatic differentiation with growth factors, we adopted a protocol from the literature (Takebe, Sekine et al. 2013). Briefly, on day 6 of culture after gut tube induction, bone morphogenetic protein 4 (BMP4, 10 ng/ml), and fibroblast growth factor 2 (FGF2, 20 ng/ml) were added from days 7-10, and hepatocyte growth factor (HGF, 10 ng/ml), oncostatin (20 ng/ml), and dexamethasone (100 nM) were added from days 10-14. Cells were assayed by qRT-PCR and immunostaining on day 14.

### RNA isolation, reverse transcription (RT) and quantitative polymerase chain reaction (PCR)

Total RNA was purified with 1) Aurum Total Mini Kit (Bio-Rad) using the spin column method with DNase 1 (Bio-Rad, Hercules, CA) (reconstituted in 10 mM tris) treatment, or 2) Trizol (Invitrogen) following manufacturer instructions. Cells were trypsinized, centrifuge pelleted, and lysed in Trizol. Chloroform was added to the lysates to separate RNA into an aqueous phase. Isopropyl alcohol was used to precipitate RNA, followed by a wash with 75% ethanol. RNA was resuspended in RNAase free water. RNA concentrations were determined by Nanodrop. RNA was converted to cDNA with an RT reaction using the iScript cDNA Synthesis Kit (Bio-Rad), and the mass of RNA was calculated such that 5 ng RNA per well to be run in the qPCR reaction. The RT reaction was performed using 5 minutes at 25° C, 20 minutes at 46° C, and 1 minute at 95°C. Reactions were then held either at 4° C or on ice. We performed 10 µL qPCR (3 µL primers at a concentration of 0.3 µM, 1 µL nuclease free water, 1 µL cDNA, and 5 µL supermix) reactions with iTaq Universal SYBR Green Supermix (BioRad) in a 96-well PCR plate (LPS). The qPCR reaction was done in a CFX96 Touch Real-Time PCR Detection System (BioRad). The qPCR reaction consisted of polymerase activation and DNA denaturation at 98°C for 30 seconds followed by 40 to 45 cycles of 98°C for 15 seconds for denaturation and 60° C for 60 seconds for annealing and extension. Melt curve analysis was performed at 65-95° C in 0.5°C increments at 5 seconds/step. Relative, normalized, gene expression was analyzed using the delta-delta-Ct method (Livak and Schmittgen 2001), with three duplicates per gene tested. Primer sequences are as shown purchased from Integrated DNA Technologies (Coralville, IA), Sigma Aldrich (St. Louis, MO), or Thermofisher (Waltham, MA) **(Supp. File 1).**

### Short-interfering RNA (siRNA) transfection of human stem cell-derived cells (forward transfection)

In a tissue culture-treated 24-well plate (LPS), 200,000 hPSC (ESC or iPSC cell lines) were plated for endoderm differentiation and hepatic differentiation protocol using the method above. Traditional transfection was performed on day 11. Positive control siRNA (siGAPDH), negative control (siScramble), and siRNA targeting FoxA1, and FoxA2 siRNA (Thermofisher) was used. Prior to transfection, cells were washed with 500 µL of fresh culture medium. Lipofectamine RNAiMAX reagent (2 µL) was mixed with Opti-MEM medium for a total of 50 µL, and 12 pmol siRNA (positive control, negative control, FOXA1, or FOXA2) was mixed with Opti-MEM medium for a total of 50 µL of siRNA mixture. Each mixture was incubated for 10 minutes at room temperature. The diluted RNAiMAX mixture and diluted siRNA mixture were mixed (1:1 ratio) at room temperature for an additional 5 minutes. Next, 100 µL of this mixture was added per well of a 24-well plate, mixed with 500 µL growth medium, and medium was not changed for 24 hours. Cells were harvested and assessed by qRT-PCR and Western Blot after 3 days, on day 14.

### Short-interfering RNA (siRNA) knockdown via reverse transfection in stable liver (HepG2) cells

Pooled siRNA targeting human FOXA1 and human FOXA2 was obtained from Dharmacon. 3 pmol of siRNA targeting FOXA1 and 3 pmol of siRNA FOXA2 were diluted into 100 μL of OptiMEM without serum per well in a 24-well tissue culture-treated plate and mixed gently. 1 μL of Lipofectamine RNAiMAX transfection reagent was added to each well and mixed gently. The mixtures were incubated for 15 minutes at room temperature. The diluted RNAiMAX mixture and diluted siRNA mixture were mixed (1:1 ratio) at room temperature for an additional 5 minutes. 500 μL of cells were resuspended in complete growth medium (DMEM and 10% FBS) without penicillin-streptomycin (Penn/Strep) at 100 cells/μL, seeded into each well, and placed in a 37°C incubator for 48 hours. Cells were collected and prepared for either qRT-PCR or western blot. Controls included cells maintained in transfection medium without siRNA, and cells treated with scrambled siRNA, and cells treated with a single siRNA pool but not both.

### shRNA sequences

Four shRNA sequences were tested and evaluated for their ability to knockdown both FOXA1 and FOXA2. Each shRNA was cloned into the host pLKO.1-puro vector. PLKO-shFOXA1#1 (Cat. #: 70095) and PLKO-shFOXA1#2 (Cat. #: 70096) were obtained from Addgene, and previously used ^45^. shFOXA2#1 and shFOXA2#2 were a kind gift from Drs. Tang and Song (Shanghai Institutes for Biological Sciences, Chinese Academy of Sciences, Shanghai 200031, China) and have been previously published. ^46^

The sequences were:

pLKO-shFOXA1#1: 5’ – TCTAGTTTGTGGAGGGTTATT – 3’;

pLKO-shFOXA1#2: 5’ – GCGTAC TACCAAGGTGTGTAT – 3’;

pLKO-shFOXA2#1: 5’ – GCAAGG GAGAAGAAATCCA – 3’;

pLKO-shFOXA2#2: 5’ – CTACTCGTACATCTCGCTC – 3’.

### shRNA lentiviral generation and transduction

Third generation lentivirus was produced with the help of the Lentivirus core facilities at Roswell Park Cancer Institute, using the cloned pLKO.1 puro vector. Transduction at multiplicity of infection (MOI) 5-10 (titer of 10^7^ particles per ml) was performed serially with shFOXA1 transduced first followed by shFOXA2 to engineer shFOXA1/2 cell lines. We performed cloning of scrambled shRNA and two shRNA sequences targeting FOXA1 and FOXA2, respectively. We engineered 4 separate lentiviruses bearing 2 separate shRNA sequences for FOXA1 and 2 separate shRNA sequences for FOXA2, for a total of 4 viruses (FOXA1 #1, FOXA1 #2, FOXA2 #1, FOXA2 #2) each of which could be selected with puromycin within the lentiviral PLKO vector. In a series of experiments (data not shown), we determined that shFOXA1 #2 and shFOXA2 #1 resulted in the most downregulation. Cells were seeded at 25% confluency in a 24-well tissue culture-treated plate one day before transduction. Cells were washed once with PBS and 250 μL of OptiMEM, 20 μL of virus, and polybrene (8 μg/mL) were added to each well for a 24-well plate. The cells were incubated for 6 hours and complete growth medium without Penn/Strep was added to each well for a total volume of 1 mL for 24-well plate. The cells were incubated for 48 hours, and media was exchanged for an antibiotic selection medium composed of complete growth medium with Penn/Strep and 1.75 μg/mL of puromycin. The cells were incubated in the selection medium for 7 days with a media change after 3 days and transduction was repeated with the second lentivirus. The cells were then assayed for gene expression with qRT-PCR and for protein expression with Western Blot. The negative control was HepG2 cells maintained in the same conditions as experimental without the virus. The positive control was done using the same transduction protocol but with a pLKO-scrambled shRNA lentivirus. Positive control cells were also selected for using 1.75 μg/mL puromycin for 7 days prior to analysis.

### Engineering of stable cell lines bearing conditional shRNA vectors

To engineer conditional cell lines which can downregulate or restore FOXA2 expression, we employed the pSingle-tTS-shRNA vector in which doxycycline (Dox) activates shRNA expression. The vector was grown in Stbl3 *Escherichia coli* with ampicillin as a bacterial antibiotic resistance. For shRNA cloning, the FOXA2 #2 shRNA was inserted downstream of the tTS promoter region using the XhoI and HindIII restriction sites. For HepG2 cell transduction, cells were plated at approximately 100,000 cells/well in DMEM + 10% FBS in a 24-well plate. Cells were transfected using FugeneHD (Promega) transfection reagent with a Fugene: DNA ratio of 4.5:1. To transfect 3 wells, a total volume of 450 μL of 0.020 μg/μL plasmid solution was made up in OptiMEM (9.9 μg of plasmid in 450 μL total volume). 45 μL of Fugene reagent was then added to each well and incubated at room temperature for 15 minutes before 150 μL of the plasmid solution complex was added to each well and mixed thoroughly. After 48 hours, the medium was changed to DMEM +10 FBS +1% P/S for expansion, or for creation of a stable cell line, cells were selected with DMEM + FBS + P/S +1000 μg/ mL G418 (Sigma-Aldrich).

### shRNA knockdown in stable liver cells with conditional shRNA vectors

For both transient transfection and stable cell line creation, knockdown was induced with the addition of doxycycline (Dox) medium. The Dox medium contained a final concentration of 1-2 μg/ml, but lower doses were tested as reported. For conditional experiments, the Dox containing media was added on ((+) Dox condition) for 3 days, and was then removed for 3 days, and cells were harvested both after Dox treatment and after Dox removal, and analyzed by qRT-PCR.

### shRNA transfection of human stem cell-derived cells

In a tissue culture-treated 24-well plate (LPS), 200,000 hPSC (iPSC cell lines) were plated for endoderm differentiation using Activin (100 ng/µL) + CHIR (3 µM) on D1, and Activin (100 ng/µL) D2-D4. From D4-D11, differentiation was carried out under SFD medium with the addition of KGF (25 ng/ml). Forward transduction methods were used on day 11, and knockdown of FOXA1 and FOXA 2 were performed by double transduction. Negative control (shScramble), shRNA targeting FoxA1, and FOXA2 shRNA was used. Lentivirus were produced by Roswell Park Cancer Institute using the pLKO.1 Puro vector. This lentivirus construction was grown in Stbl3 with 100 ug/mL of ampicillin as the vector expressed ampicillin resistance gene. This shRNA was cloned in between AgeI and EcoRI restriction enzyme cut sites. On day 11 of differentiation, 100 µL of shFOXA1, 100 µL of shFOXA2, and 300 µL of OptiMem, polybrene (8 µg/mL) was added per well. The virus titer is of (1 x 10^7^ IU/ml) and was used at an MOI of 5. After 6 hours of incubation in transduction complex, 250 µL of SFD medium was added w/o addition of P/S. Media changes was then performed every 24 hours w/o addition of P/S. Cells were harvested 3 days post-transfection for qRT-PCR or Western Blotting.

### Immunofluorescence

Cells were seeded at 50,000 cells/well in a 24-well tissue culture-treated plate one day before staining. Cells were rinsed three times with PBS at a volume of 500 µL per well. All wash steps were performed with 500 µL PBS/well unless otherwise specified. Cells were fixed with 500 µL of 4% paraformaldehyde at room temperature under cell culture hood. Cells were washed again three times for 5 minutes each wash before incubating with 1% PBS-Triton X for 30-60 minutes for permeabilization. Cells were rinsed thrice more before incubation in 1% BSA/PBS (Blocking Buffer) for 1 hour. Primary antibody incubation was done overnight at 4°C at a 1:500 dilution in either blocking buffer or PBS. After overnight incubation, cells were washed 4 times for 15 minutes before 1-hour incubation with a fluorescently labeled secondary antibody, diluted 1:500 in either blocking buffer or PBS. After secondary antibody incubation, cells were washed 4 times for 15 minutes each using 500 µL of 0.1% PBS-Triton X, cells were rinsed once more with Milli-Q water before adding 500 µL of a 1:1000 Dilution of DAPI in PBS before imaging. Cells were Imaged with Zeiss Axiovision SE64.

### Protein isolation and Bradford assay

On the day of protein collection, cells were washed once with PBS and then incubated in 300 μL of 0.05% Trypsin for 6-10 minutes. 500 μL of DMEM containing 1% v/v of Pen-Strep (P/S) and 10% fetal bovine serum (FBS) was added and cells were then collected and transferred to sterile microfuge tubes. Cells were then pelleted at 5,000 RPM for 30 seconds and cell pellet was resuspended in 300 μL of 1x PBS, in which cells were washed twice. For the last wash, cell pellets were resuspended in 250 μL of RIPA buffer containing Protease inhibitor and incubated on ice for 5 minutes. Cells were then spun at 4°C for 15 minutes at 14,000 RPM and the supernatant was collected for assaying via Bradford assay. Bradford assays were conducted by diluting protein samples by a factor of 10 with Milli water, and 5μL of each sample or known standard was platted into each appropriate microplate well of a 96-well plate. 250 μL of Coomassie protein assay reagent (ThermoScientific) was added to each well and was left to mix on a plate shaker for 30 seconds. Absorbances were measured at 595 nm with a plate reader and a standard cure was constructed using the average of triplicated samples and known protein concentrations, a best fit line constructed using a 3rd degree polynomial was used to calculate the unknown sample concentrations.

### Western blot and quantitation

Equal concentrations of protein were loaded in all lanes of a 10% polyacrylamide gel with a 4% stacking gels were used and prepared per manufacturer’s instructions (Invitrogen). Pre-made, Bolt 4-12% Bis-Tris Plus 10-well gels (Invitrogen) were also used. Protein sample preparation involved using equal concentrations of 10-50 μg/well for a final volume of 25 μL, for use in a 10-well polyacrylamide gel. Samples were heated, loaded, and run in 1X SDS-PAGE buffer for 45 minutes at 125V (Bio-Rad Mini Gel Tank). Pre-made gels were run at 200V for 22 minutes. Next, gels were washed and placed in a gel transfer assembly (Mini Blot Module, Invitrogen) for transfer to a nitrocellulose membrane (Bio-Rad) for approximately 60 minutes. Some transfers took place using the iBlot2 transfer module (Invitrogen) using nitrocellulose transfer stacks (Invitrogen). The membrane was washed in MilliQ water 2 times for 5 minutes each before being placed in 15 mL of blocking buffer consisting of 5% milk powder in PBS-Tween (PBS pH 7.4 + 0.1% Tween 20) for 1 hour at room temperature, washed, and incubated overnight with primary antibody overnight, washed three times, and incubated for 1 hour at room temperature in HRP-conjugated secondary antibodies, which were diluted in blocking buffer at a 1:100-1:5000 dilution. After incubation in secondary antibody, membranes were washed again in PBS-Tween before being placed in 0.1 mL working solution (equal parts stable peroxide solution and Luminol/Enhancer solution) per cm^2^ membrane for 5 minutes before being placed for imaging in ChemiDoc Gel Imaging System (Bio-Rad). Gels were quantified using image J by taking the ratio of the integrated intensity (area x intensity) at for a specific gene and normalizing it to the integrated intensity of the house keeping gene.

### Oil-Red-O assay

Oil-Red-O (Sigma Aldrich) stock was prepared by dissolving 300 mg of Oil-Red-O in 99% isopropanol. 50,000 control (scrambled) and shFOXA1/2 knockdown cells were seeded on Day 0 in 24-well plate until the end of Day 4. Culture medium (High glucose DMEM + 10% FBS+ 1% Pen/Strep) was changed every other day. 1.75 µg/ml puromycin was added to the medium for shFOXA1/2 knockdown cells. On Day 4, cells were fixed by washing with PBS and incubation in 10% formalin for 1 hour at RT. Then, working solution was made by mixing stock with sterile water (3:2) and sterile filtration (0.22 µm filter) in the dark. After formalin fixation, cells were incubated in 60% isopropanol for 5 min. Working solution was added and incubated for 5 min shaker at room temperature. Cells were rinsed with sterile water until the solution runs clear. Next, hematoxylin was added, and cells were incubated for 1 min. Cells were then rinsed with warm sterile water until the solution runs clear. Cells were then imaged by brightfield and phase microscopy. Images were analyzed by ImageJ by dividing the area of Oil-Red-O by area of cells.

### Microscopy

Cell lines and stem cell cultures were imaged with a benchtop microscope (EVOS fluorescent, phase contrast microscope, #AMEFC4300R) at 4x, 10x, and 20x or with the Zeiss Axiovision SE64 microscope. Images were acquired and stored and used to visualize cells.

### RNA-sequencing (RNA-seq)

Total RNA integrity was determined with the Agilent Technologies Fragment Analyzer RNA assay system. The TruSeq Total RNA Stranded Library Preparation kit (Illumina) with rRNA removal was used for preparation of RNA sequencing libraries. Multiplexed libraries were individually quality controlled with the Fragment Analyzer and quantified using the Quant-iT ds DNA Assay Kit (Invitrogen). The libraries were pooled to 10 nM and diluted for qPCR using the KAPA Library Quantification Kit (Kapa Biosystems). Subsequently the pooled libraries were normalized to 4 nM based on qPCR values. All samples were sequenced on the NextSeq500 (Illumina) in midoutput mode producing 160 million paired end 75 bp reads, ∼25 million per sample.

### RNA-seq (Transcriptome) analysis

The raw fastq files were first aligned using the hisat2-2.1.0 GRCh38 genome and converted into sam files. The sam files were converted into bam files using samtools-1.6. FeatureCounts was used to count reads for genes from the GRCh38 genome to create a text file with the two controls and three experimental samples. The text file was imported into R where it was analyzed with DESeq2. The gene expression data was normalized with Rlog normalization in DESeq2. The built in prcomp function was used on the normalized data to find the first and second principal component analysis (PCA1 and PCA2). PCA1 and PCA2 were plotted using R plotting tools. The volcano plot figure was created by importing the same count data into R. The DESeq2 function lfc shrink was used to shrink the data using the contrast between the condition and experimental groups. The res function was used to obtain the adjusted p-value and log2-fold change for each gene. The log2-fold change was then plotted on the x axis and the -log10 of the adjusted p-value was plotted on the y axis. The genes were colored if the adjusted p-value was under 0.05. The genes in red represent upregulation and the genes in blue represent down regulation based on the log2-fold change. The heatmap was created with the gplots function heatmap.2 in R. The normalized number of reads for each gene were compared between samples to determine the color. Green represents relatively higher gene expression, while red represents relatively lower gene expression. The 683 genes with an adjusted p-value under 0.05 were plotted and divided into upregulating and downregulating groups and listed by increasing adjusted p-value for each group. Every seventh gene in the list was labeled. Unbiased hierarchical clustering was used to group the samples with the built in hclust function within heatmap.2. The 683 significantly expressed genes were additionally analyzed with Gene Ontology Panther classification system using the “Statistical Overrepresentation Test” to find significant biological processes, cellular components, and molecular functions. Default options were used in this analysis except the reference list was replaced by a cumulative gene list including all genes found to have significant expression (adjusted p-value<0.05) from the RNA-seq analysis. Significance was determined by a (FDR)-qval <0.05.

### RNA-seq normalized enrichment score (NES) analysis

Normalized enrichment scores (NES) for differential gene sets were analyzed using Gene Set Enrichment Analysis (GSEA). Gene expression data was normalized with the median of ratios method in DESeq2. The normalized read counts of the genes were compared between the experimental group (shFOXA1/2 -/-) and control group with a Student’s t-test method and the genes were arranged from most upregulated to most downregulated. The GSEA 4.0.3 software from the Broad Institute analyzed the list using the GSEAPreranked tool. All default settings were used except the chip platform field was set to “no collapse” and the max set size was set to 2000. The version 7.0 gene sets from Gene Ontology (GO) and KEGG were downloaded from the Broad Institute and used as the gene sets database. The results were filtered for p-value<0.05, NES -1.5 > or < 1.5, and either (FDR)-qval or (FWER)-pval<0.1. The results were grouped based on enrichment and were sorted based on “SIZE.” The signal-to-noise method was also used to sort the genes instead of the Student’s t-test and obtained similar results.

### Identification of liver GRN promoter/enhancer binding sites

To determine if genes downregulated in the shFOXA1/2 condition were likely to be regulated by Foxa, HNF4α, or HNF1, we analyzed a candidate list of hepatic genes that were downregulated. Information available in the “Genomics” section for each gene of interest through Genecards Suite (<u>genecards.org)</u> was used to identify transcription factor binding sites in the promoter/enhancer sequences for the gene. We also tested the ENCODE database which was a less exhaustive list of promoter binding sites (https://genome.ucsc.edu/ENCODE/) and corroborated the list that was generated using Genecards ^47^. Genecards uses a technique called Genehancer, which utilizes the ENCODE database, as well as 3 other databases, and it is therefore broader. Listed in **Table 3** are the liver GRN (Foxa, HNF4α, HNF1ß) and the non-liver GRN. The non-liver GRN column does contain TF binding sites for some liver GRN, but we have separated it this way based on our current analytic approach.

### Assessment metabolic genes of shFOXA1/2 cells

GSEA metabolism gene sets from Kegg, BioCarta, and Reactome were further studied and visualized. The genes making up these gene sets were assessed using DESeq2 based on log-2-fold-change and p-value. A gene was determined to be significant if for P < 0.05 and a log2-fold-change > 0.5 or < -0.5. Trending genes for metabolic gene analysis were defined as P < 0.20, due to the small sample size of the analysis. These trending genes are shown with P values in text or in tables.

### Functional metabolic analysis of shFOXA1/2 cells

Cell energetics were assessed with a Seahorse XFe24 Extracellular Flux Analyzer (Seahorse Bioscience). Control or shFOXA1/2 cells were plated at a density of 40,000 cells/well into XFe24 microplates and cultured at 37 °C with 5% CO_2_ overnight. Cell culture medium was replaced with either Seahorse XF Base Medium supplemented with 2 mM L-glutamine for the glycolysis stress test or Seahorse XF Base Medium supplemented with 2 mM L-glutamine, 1 mM pyruvate, and 10 mM glucose for the mitochondrial stress test. Extracellular acidification rate (ECAR) was determined using the Seahorse XF Cell Glycolysis Stress Test following the manufacturer’s instructions. Injections consisted of glucose (10 mM), followed by oligomycin (1 µM), followed by 2-deoxyglucose (50 mM) in Seahorse glycolysis stress test media. Three measurements were taken prior to the first injection and after each injection. Oxygen consumption rate (OCR) was assessed using the Seahorse XF Cell Mito Stress Test according to the manufacturer’s instructions. Injections consisted of oligomycin (1 µM), followed by carbonyl cyanide-4 (trifluoromethoxy) phenylhydrazone (FCCP) (2 µM), and then rotenone (0.5 µM) in Seahorse mitochondria stress test media. Three measurements were taken prior to the first injection and after each injection.

### Urea Assay

QuantiChrom Urea Assay Kit (DIUR-100) was purchased from BioAssay Systems, stored at 4°C, and the manufacturer instructions were used to measure urea concentration within the medium produced by hepatic cells. Control (HepG2) and shFOXA1/2 cells were plated at 200000 cells/well in 24-well plates. Medium was collected after 24 hours and stored at 4°C overnight prior to analysis. All reagents and samples were placed at room temperature. Fresh working reagent was prepared by combining Reagent A and Reagent B. The standard solution containing urea was serially diluted between 1.4 mg/dl and 50 ng/ml. In a 96-well plate, 50µl of sample medium, (n = 3), blank (medium), and standards were added to the wells, and each condition was performed in triplicate. 200 µl of fresh working reagent was added to each well and the plate was mixed. The plate was incubated 50 minutes. The absorbance (optical density) was read at 430 nm on the spectrophotometer (Biotek Synergy 4-plate reader). A standard curve of the absorbance of the serially diluted standards from 1.4 mg/ml to 50 ng/ml was used to calculate the sample concentrations. Standards were diluted as needed. Urea concentration was calculated by subtracting the absorbance of sample from blank, the absorbance of the standard by the blank, and dividing these two number by each other.

### Statistics

For statistical comparison between two groups, Student’s t-test was performed, and p-values equal to or less than 0.05 was considered to have statistical significance and are delineated within the text and figures/figure legends. Statistics for the RNA-seq analysis is detailed in those sections.

## RESULTS

### FOXA 1/2 regulates the endoderm and liver GRN, and liver differentiation genes, in a stable liver cell line

We hypothesized that siRNAs targeting both FOXA1/2 regulate ALB expression, the GRN in human liver cells, and lead to loss of liver differentiation. We chose to apply this approach to human HepG2 cells, a well-established model of hepatocyte gene regulation ^48–49^. We first established the siFOXA1/2 phenotype in HepG2 cells. We performed single knockdown studies siRNA targeted only to FOXA2 or to FOXA1/2, and found partial knockdown of the target TF, but not the corresponding FOXA factor (data not shown). This suggested that FOXA1 compensates for FOXA2, and vice versa, as previously reported in murine systems ^15^. We also performed dose response studies, and we found 3 pmol of siRNA for both FOXA1 and FOXA2 (total of 6 pmol) resulted in optimal knockdown. Knockdown did not affect morphology or proliferation **(Fig. 1A)**. qRT-PCR analysis demonstrated that knockdown of FOXA1/2 resulted in a downregulation of FOXA1, FOXA2, and ALB, but not AFP **(Fig. 1B)**. Because of decreased ALB, we hypothesized there was de-differentiation, and we tested master factors (stem cell markers) of alternate lineages **(Fig. 1C)**. We found statistically significant upregulation of NANOG (pluripotent), PAX6 (neuroectoderm), SOX7 (visceral endoderm), and CXCR4 (endoderm), in the siFOXA1/2 knockdown cells **(Fig. 1C)**. Decreased ALB was possibly due to decreased hepatic GRN, and we observed a significant downregulation in HEX, **(Fig. 1C)**, and we also identified trends in PDX1 (pancreas) and BMP4 (mesoderm) upregulation (data not shown). The downregulation of FOXA1/2, ALB, was confirmed by Western blot, while siFOXA1 results in ALB upregulation **(Fig. 1D-F)**, and by immunostaining **(Fig. 1G-H)**. However, we still did not observe any changes in several of the hepatic GRN **(Fig. 1C)**. We reasoned that a second siRNA sequence may have more potent effects on FOXA1/2 and on hepatic GRN knockdown.

**Figure 1.**
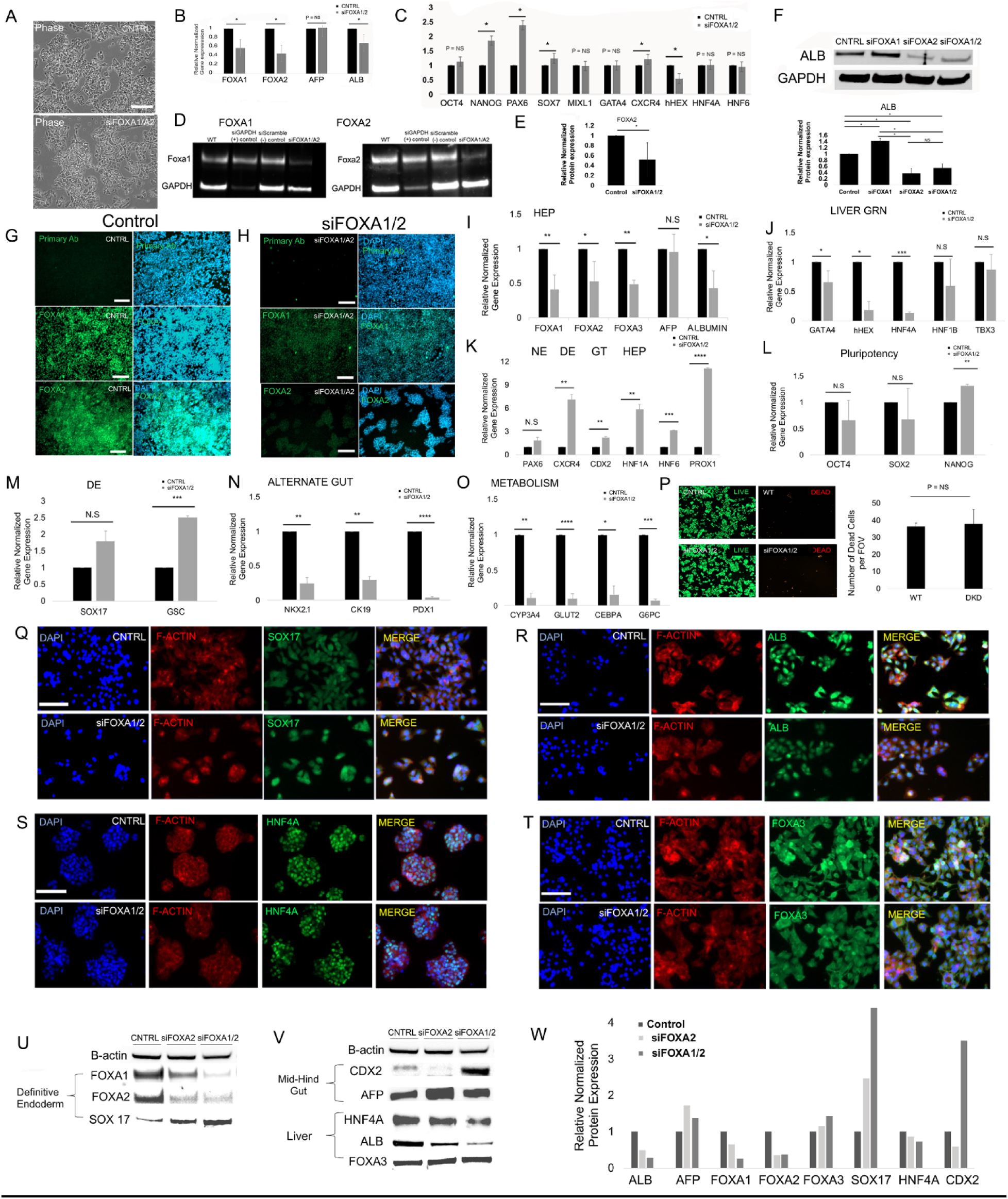
siRNA of FOXA1/2 regulates differentiation genes and GRN TF in a liver cell line (HepG2) A) Above-Phase contrast images for morphological comparison of scrambled (top) and siRNA FOXA1/2 48 hours after transfection in HepG2 cells. Bar = 100 µm. B) Bar graph of gene expression (qRT-PCR) data for siFOXA1/2 knockdown in HepG2 assayed after 48 hours, compared to scrambled controls. N = 5 for control (scrambled) and siRNA FOXA1/2 conditions. FOXA1, P = 0.000021, FOXA2, P = 0.000014, AFP, P = NS, ALB, P = 0.0003. Plotted is mean ± SD. Significance (*) defined as P ≤ 0.05. C) Same as B) except further genes tested. Oct 4, P = NS, NANOG, P = 0.00078, PAX6, P = 0.00625, SOX7, P = NS (P = 0.102), MIXl1, P = NS, GATA4, P = NS, CXCR4, P =NS, HEX, P = 0.0036, HNF4A, P = NS, HNF6, P = NS. Plotted is mean ± SD. Significance (*) defined as P ≤ 0.05. D) Western blot of HepG2 cells 48 hours after transfection for FOXA1 (left) and FOXA2 (right). The lanes from left to right are 1) Control (HepG2), 2) siGAPDH (positive control), 3) Scrambled siRNA, 4) siFOXA1/2 (siFOXA1 and siFOXA2). E) Quantitative analysis of data in D). Significance (*) defined as P ≤ 0.05 (n = 3). F) Western Blot of ALB in control, siFOXA1, siFOXA2, and siFOXA1/2 conditions, with quantitative analysis below. Significance (*) defined as P ≤ 0.05 (n =3). G) Immunohistochemistry of FOXA1 and FOXA2, 48 hours after control (siScramble) transfection in HepG2 cells. FITC (left) and DAPI/FITC (right). Bar = 100 µm. H) Same as G) except siFOXA1/2 condition. I) Same as B) except an alternate siRNA molecule targeting FOXA1/2 in HEPG2 cells is analyzed. J) Same as I) except downregulation of the hepatic GRN shown K) Same as I) except upregulation of Neuroectoderm (N) TF, definitive endoderm (DE) marker, gut tube endoderm (GT) TF, and hepatic GRN (HEP). L) Same as I) except pluripotency TF. M) Same as I) except DE genes are shown. N) Same as I) except alternate gut genes are shown. O) Same as I) except metabolism genes are shown. P) Fluorescence images of live (green) dead (red) assay in control (scramble) and shFOXA1/2 conditions Q) Immunofluorescence of CNTRL (Scrambled) and siFOXA1/2 for (DAPI, F-ACTIN, SOX17) in HepG2. R) Same as M except immunostaining for ALB S) Same as M except immunostaining for HNF4A T) Same as M except immunostaining for FOXA3 U) Western blot for endoderm TF (FOXA1/2, SOX 17) same experiments as in I)-P) with alternate siRNA targeting FOXA1/2 in HEPG2 cells V) Same as U) except gut tube and liver markers and liver GRN. W) Quantitation of blot is R)

Therefore, we tested another set of commercially available siRNA sequences, and observed a more de-differentiated phenotype. We again demonstrated knockdown of FOXA1/2, and downregulation of ALB in the siFOXA1/2 phenotype **(Fig. 1I)**. When we tested liver GRN, we observed significant knockdown of not only hHEX, but also HNF4A and GATA4, with HNF1B and TBX3 trending downwards **(Fig. 1J)**. Moreover, we observed significant upregulation of PAX6 (neuroectoderm), CXCR4 (endoderm), CDX2 (gut tube), HNF1A(hepato-biliary), HNF6 (hepato-biliary), PROX1(hepatic) markers **(Fig. 1K)**. Further, we observed a significant upregulation of NANOG (pluripotency), and trending downregulation of OCT4 and SOX2 **(Fig. 1L)**. We also found upregulated DE TF including GSC (significant) and SOX17 (trending) **(Fig. 1M)**. We found that gut derivative markers NKX2.1 (lung), CK19 (Biliary), and PDX1 (pancreatic) were significantly downregulated **(Fig. 1N)**. Further, we found that metabolic liver markers CYP3A4, GLUT2, CEBPA, and G6PC were significantly downregulated **(Fig. 1O)**. We also confirmed that there no change in viability in the siFOXA1/2 phenotype **(Fig. 1P)**. Overall, this second siRNA sequence resulted in a more extensive knockdown phenotype. Immunohistochemistry of SOX17 (endoderm) confirmed upregulation, and ALB, which confirmed downregulation **(Fig. 1Q-R)**. Immunohistochemistry of HNF4A and FOXA3 demonstrated no major differences **(Fig. 1S-T)**. Western blot confirmed significant knockdown of FOXA2 and confirmed dedifferentiation with expected expression of TFs and differentiation markers. The data with the new siRNA sequence confirmed previous findings, including the downregulation of FOXA1, FOXA2, and ALB **(Fig. 1U)**. Further, we observed an upregulation of DE marker SOX 17, and CDX2, showing de-differentiation **(Fig. 1V)**. We also observed downregulation of HNF4A **(Fig. 1V-W)**. These data demonstrate siFOXA1/2 demonstrates loss of hepatic differentiation, and gain of pluripotent, neuroectoderm, endoderm, gut tube, and early liver (HNF1, HNF6, PROX1) markers.

### The shFOXA1/2 phenotype demonstrates a downregulation of liver differentiation genes and TF that comprise liver GRN

We wanted to further confirm this siFOXA1/2 phenotype. We employed multiple, published, viral sequences, and we determined the sequences we labeled shFOXA1 #2 and shFOXA2 #1 resulted in the most downregulation. We determined that serial transduction of shFOXA1 + shFOXA2, followed by antibiotic selection, resulted in a stable shRNA FOXA1/2 phenotype. Optimization experiments were performed **(Fig. 2A)**, confirming the most potent sequences, shFOXA1 #2 and shFOXA2 #1 viruses, as shown by a representative (n = 1) transfection experiment. We then determined the shFOXA1/2 phenotype. We again analyzed gene expression of core TF and differentiation markers including pluripotency (Nanog), neuroectoderm, (PAX6), endoderm (GSC, CXCR4), pancreas (PDX1, SOX 9), biliary (CK19), and the endoderm/hepatic nuclear GRN TFs (FOXA3, HNF4A, HNF1B, HEX, GATA4, TBX3). The data demonstrated a significant decrease in ALB **(Fig. 2B)**, and trending increases in AFP, CXCR4, and CK19. We also observed significant upregulation in GSC (endoderm) expression (1.5-fold), PDX1 downregulation a (0.84-fold), and a trending increase in PAX6 and decrease in SOX9 expression **(Fig. 2C)**. The shRNA FOXA1/2 phenotype showed significant changes in core members of the endoderm/gut tube/liver GRN, including HNF4A, HNF1B, HEX, and TBX3 **(Fig. 2D)**, and upregulation in FOXA3 and GATA4 **(Fig. 2D)**. Thus, there was a strong downregulation of liver differentiation genes and core liver GRN that establishes a liver cell fate, with a concomitant increase in early liver markers like FOXA3, AFP, and CXCR4. Further, PAX6 (neuroectoderm), and CK19 (biliary) were significantly increased. Western blotting demonstrated that shFOXA1/2 downregulated FOXA2 more than shFOXA1 or shFOXA2, suggesting it has the strongest phenotype, and this downregulation was significant **(Fig. 2E)**. Western blotting confirmed significant downregulation of FOXA1, ALB, HNF4A, TBX3, and upregulation of AFP and CDX2 **(Fig. 2F)**. These data demonstrate a coordinated decrease in liver differentiation, and evidence of immature states (hepatic endoderm) and alternate fates (biliary, neuroectoderm) consistent with a de-differentiated state, similar to the siRNA phenotype **(Fig. 1)**.

**Figure 2.**
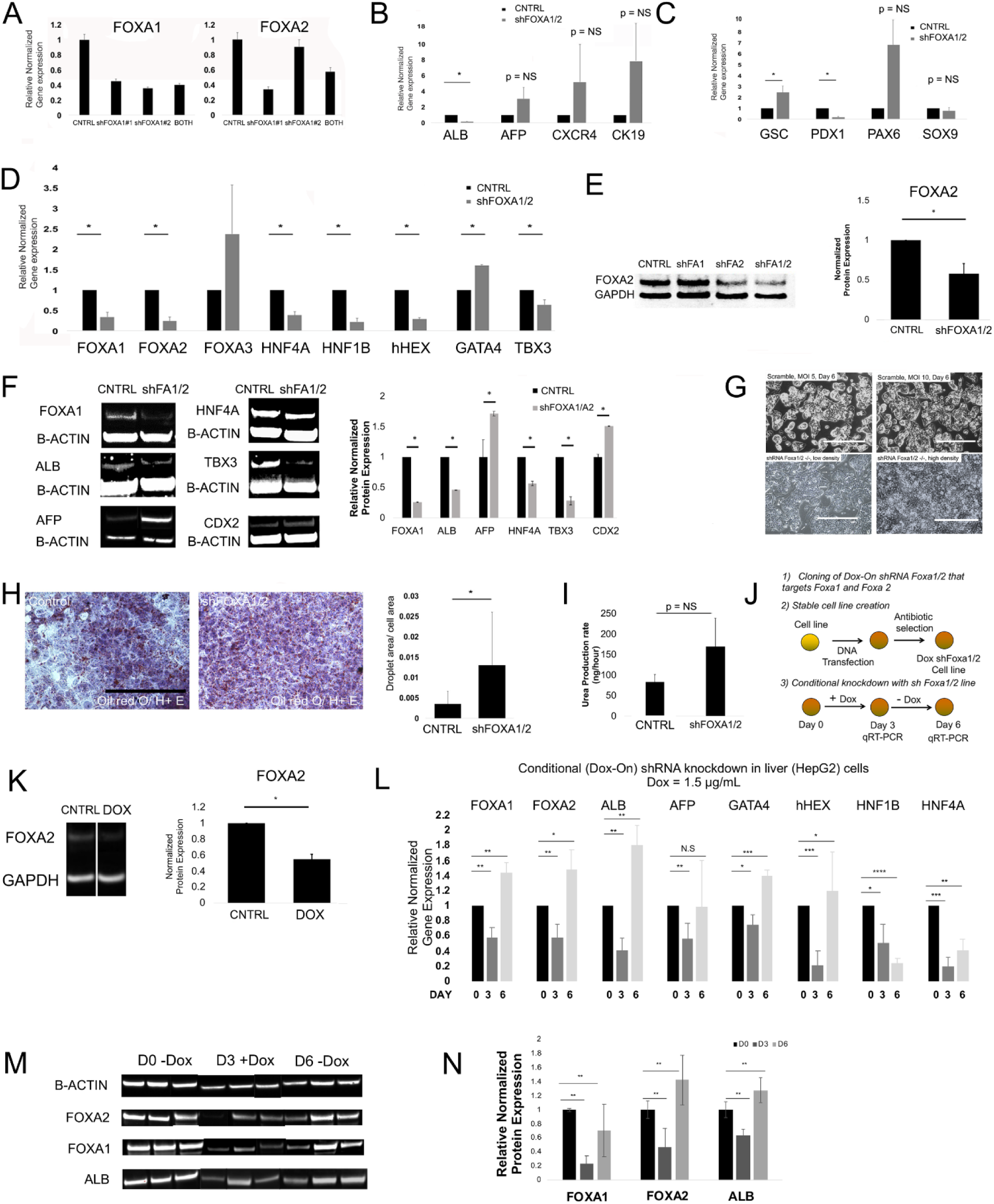
shRNA FOXA1/2 knockdown regulates liver differentiation, GRN TFs, and function. A) Bar graph of gene expression (qRT-PCR) after shRNA transduction with shFOXA1, shFOXA2, shFOXA1/2, and scrambled shRNA control, followed by antibiotic selection. N = 1 shown. Left - FOXA1 expression in control, shFOXA1, shFOXA2, and both shFOXA1/2. Right-FOXA2 expression in control, shFOXA1, shFOXA2, and both shFOXA1/2. Plotted is, mean ± SD for 3 replicates. B) Bar graph of gene expression (qRT-PCR) after shRNA transduction with shFOXA1/2, and scrambled shRNA control, followed by antibiotic selection for two weeks. ALB, P = 0.000019, AFP, P = NS, CXCR4, P = NS, CK19, P = NS. Plotted is mean ± SD. Significance (*) defined as P ≤ 0.05 (n = 3). C) Same as B) except N = 3 for all conditions. GSC, P = 0.027, PDX1, P = 0.0002, PAX6, P = NS, SOX9, P = NS. Plotted is mean ± SD. Significance (*) defined as P ≤ 0.05 (n = 3). D) Bar graph of gene expression (qRT-PCR) after shRNA transduction of HepG2 cells with shFOXA1/2, and scrambled shRNA control, followed by antibiotic selection for two weeks. FOXA1, P = 0.001, FOXA2, P = 0.004, FOXA3, P = NS, HNF4α, P = 0.0006, HNF1ß, P = 0.0002, HEX, P = 0.000008, GATA4, P = 0.0000009, TBX3, P = 0.013. Plotted is mean ± SD. Significance (*) defined as P ≤ 0.05 (n = 3). E) Western blotting for scrambled shRNA (control), shFOXA1, shFOXA2, shFOXA1/2 conditions. FOXA2 is shown. Quantitation of Western blot (control and shFOXA1/2 condition) demonstrates P = 0.00007 (n =5). F) Western blotting for scrambled shRNA (control) and shFOXA1/2 conditions. FOXA1, ALB, AFP, HNF4A, TBX3, and CDX2 shown. Quantitation of Western blot demonstrates FOXA1, P = 0.000002, ALB, P = 0.000048, AFP, P = 0.0014, HNF4α, P = 0.015, TBX3, P = 0.017, and CDX2, P =.0.00005 (n =5). G) Morphology comparison between transfected and non-transfected HepG2 cells-Top row-shRNA scramble at MOI of 5 and MOI of 10, Bottom row-shRNA FOXA1/2 knockdown condition. Bottom left panel-Small white arrows demonstrate elongated and flattened cells at the edges of colonies, that were seen routinely at low density. Bottom right panel-shFox2 knockdown cell lines at high density. (Bar = 250 µm). H) Brightfield color images of control and shFOXA1/2 cells stained with Oil-red-O and counterstained with Hematoxylin and Eosin. Quantitation of staining to the right. Plotted is mean ± SD. Significance (*) defined as P ≤ 0.05 (n = 3). Bar graph of urea production rate (ng/h) between control and shFOXA1/2 conditions. Plotted is mean ± SD. P = NS. (n = 3). J) Schematic of stable cell line creation for conditional shFOXA1/2 phenotype. Cells were transfected with Doxycycline (Dox)-On shRNA vector (shRNA FOXA2#1) targeting both shRNA for FOXA1 and FOXA2. (1) shRNA was cloned into the Dox-on vector. (2) Next, DNA was transfected and selected for by antibiotic or shRNA scrambled controls, continuously selected after several passages with puramycin selection, and analyzed. (3) Next, cells are exposed to Dox (1.5 μg/ml) for 3 days and then Dox is removed for 3 days, with cell collection at Day 3 and Day 6 for qRT-PCR. K) Western blot of control and Dox conditions (1.5 μg/ml). Bar graph of Western blot analysis with FOXA2 expression in control conditional and shFOXA1/2 cell line. Significance (*) defined as P ≤ 0.05 (n = 3). L) Bar graph of gene expression (qRT-PCR) in conditional shFOXA1/2 cell line on Day 0, Day 3 (Dox addition, from Day 0-Day 3), Day 6. (Dox removed from Day 3-Day 6). Dox concentration was (1.5 μg/ml) Significance defined as P < 0.05, **, P <0. 01, ***, P < 0.001 (n = 3). M) Western blot of FOXA2 expression in conditional Dox-On shFOXA1/2 cell line, demonstrating knockdown and upregulation, indicating reversibility. Day 0 (-Dox), Day 3 (+ Dox), Day 6 (-Dox). N) Quantitation of Western Blot in M). Significance defined as *, P < 0.05, **, P <0. 01, ***, P < 0.001. (n = 3).

Our shFOXA1/2 double knockdown cell line demonstrated no change in morphology **(Fig. 2G, top row)**, although at colony edges we observed flat, elongated cells **(Fig. 2G, bottom row)**. We also observed changes in refractile vesicles in shFOXA1/2 cells, which we determined were potentially oil droplets. We performed Oil-Red O staining, which demonstrated a significance increase in intracellular lipid **(Fig. 2H)**, consistent with the role of FOXA1 in lipid metabolism ^50^. To evaluate differences in liver-specific function, we performed urea analysis and demonstrated urea synthesis was trending upwards in shFOXA1/2 cells (n = 3, P = NS (p = 0.1)) **(Fig. 2I)**. Overall, shFOXA1/2 cells exhibited hepatic de-differentiation, downregulation of GRN TF, upregulation of markers/GRN TF for alternate cell fates, and increased intracellular lipid. To test another model of shRNA downregulation, we engineered a DNA transfected (nonviral), Doxycyline (Dox)-on shFOXA1/2 cell line that expresses the Dox-on, pSingle-tTS-shRNA vector, in which shRNA for FOXA1/2 was controlled by the addition of Dox. After cloning the shFOXA2 #2 shRNA and scrambled shRNA into the pSingle vector, we engineered stable cell lines selected by G418 antibiotic **(Fig. 2J)**. We first determined that 1.5-1.75 ng/ml was optimal for knockdown response and no toxicity. Dox addition significantly knocked down FOXA1/2 **(Fig. 2K)**. We then tested reversibility of FOXA1/2. We performed addition of Dox (1.5 ng/ml) for 3 days, followed by removal for an additional 3 days, followed by analysis **(Fig. 2L)**. The Dox-sensitive shFOXA1/2 line demonstrated similar morphologic features to the shFOXA1/2 line **(Fig. 2K)**. With Dox addition, we observed downregulation of FOXA1, FOXA2, ALB, AFP, GATA4, HEX, HNF1B, and HNF4A **(Fig. 2K)**. After Dox removal, we observed upregulation of FOXA1, FOXA2, ALB, GATA4, HEX, a more moderate upregulation of HNF4A, and a continued downregulation of AFP and HNF1ß **(Fig. 2K)**. Importantly, FOXA1, FOXA2, ALB, GATA4, hHEX on day 6 were significantly higher than the day 0 levels. Western blotting for FOXA1, FOXA2, and ALB, demonstrated a downregulation and upregulation, as expected **(Fig. 2M-N)**. Thus, the global changes induced by Dox were reversible, but restoration of the same value as previous did not occur.

### Global transcriptome analysis of shFOXA1/2 knockdown phenotype in liver cells demonstrates increased stem cell/ cell differentiation genes of alternate fates, and downregulation of metabolic genes

We hypothesized that the shRNA phenotype may result in global, genome-wide effects. We performed whole transcriptome analysis on control (n = 2) and experimental (n = 3) conditions. Principal component analysis (PCA) figure shows significant clustering of samples **(Fig. 3A)**. The volcano plot **(Fig. 3B)**, demonstrates 448 upregulated genes (green) and 235 downregulated genes (red) between the two conditions (**Supp. File 2**), for a total of 683 genes. In our heat map analysis, using unsupervised, hierarchical clustering, we found PAX6 in the upregulated gene set, and ALB in the downregulated gene set, consistent with qRT-PCR data **(Fig. 3C)**. A full list of the differentially regulated genes is provided **(Supp. File 2)**, with further analysis **(Supp. File 3**, Tables 1-2). On this list included genes involved in axon guidance and vascular development (NRP2, neuropillin 2), kidney epithelial function (Cdh16, cadherin 16), kidney cell interactions (SYNPO, synaptopodin), and cardiac development (Alpk2 (alpha kinase 2)). Interestingly, these alternate lineages have been shown in meta-analysis of human stem cell-directed hepatic differentiation studies ^51^. Among the top 30 downregulated genes include those involved in energy metabolism (PPARGC1A (PPARG coactivator 1 alpha, PGC1α)), liver development and liver differentiation and function (ALB), and drug and steroid metabolism (AKRC12 (Aldo-Keto Reductase Family 1 Member C2)) (**Table 2**). We analyzed all the normalized RNA-seq data, not just differentially regulated genes using the Gene Set Enrichment Analysis (GSEA) software to find significantly enriched gene sets based on normalized enrichment score (NES) and false discovery rate (FDR) **(Fig. 3D, Supp. File 4)**. We also observed over 500 genes associated with epithelial cell differentiation, followed by processes associated with skin development, striated muscle differentiation, eye development and neuron migration **(Fig. 3D, Supp. Files 4-5)**. We observed processes with downregulated genes, including stem cell differentiation, anterior-posterior specification, **(Fig. 3D, Supp. File 4)**. Overall, this suggests that shFOXA1/2 knockdown resulted in a downregulation of liver GRN, and in a global block liver differentiation, which was concomitant with an upregulation in differentiation markers in other major lineages like mesoderm and neuroectoderm. Interestingly, similar lineages (cardiac, neural) are upregulated during human stem cell hepatic differentiation ^51^, FOXA1/2/3 knockdown in liver tissues ^16^, and FOXA1/2 knockout in pancreatic islets ^52^. To further validate our transcriptome data, we hand-picked a series of downregulated hepatic genes of interest, and examined whether their regulatory regions had a FOXA1/2, HNF4α, or HNF1 binding site. Nearly all genes had at least one of these binding sites (**Table 3**). This data provided a further link between knockdown of FOXA1/2, loss of HNF4a and HNF1, and downregulation of several liver-specific genes.

**TABLE 1.**
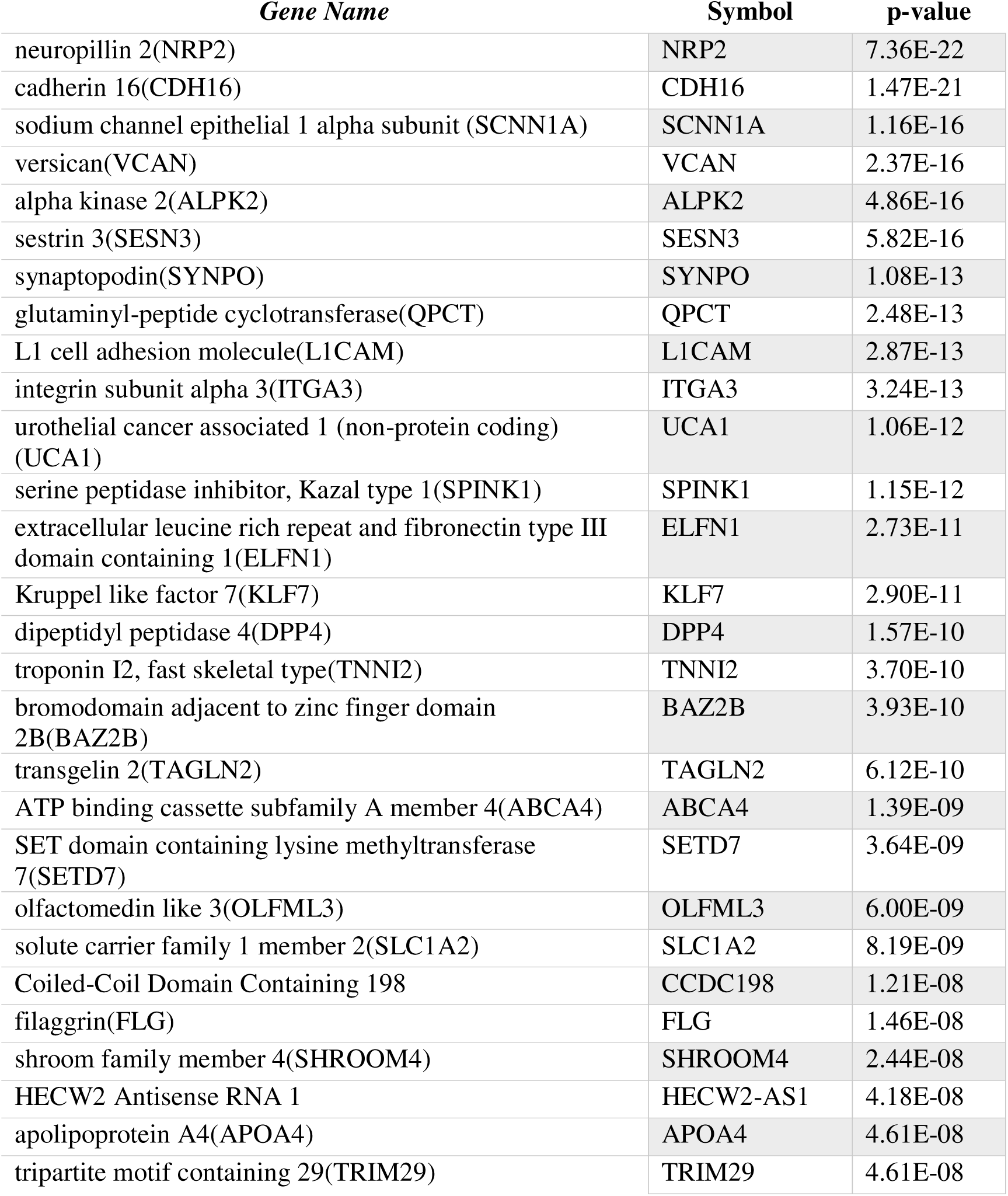
Top 30 Upregulated genes in shFOXA1/2 knockdown cells compared to HepG2

**TABLE 2.** Top 30 Downregulated genes in shFOXA1/2 knockdown cells compared to HepG2

| *Gene Name* | *Symbol* | *p-value* |
| --- | --- | --- |
| metallothionein 1G | MT1G | 1.32E-12 |
| aldo-keto reductase family 1 member B10 | AKR1B10 | 5.16E-12 |
| metallothionein 1E | MT1E | 7.51E-11 |
| aldo-keto reductase family 1 member C1 | AKR1C1 | 2.13E-09 |
| alpha-1,6-mannosylglycoprotein 6-beta-N-acetylglucosaminyltransferase | MGAT5 | 2.17E-09 |
| cadherin like and PC-esterase domain containing 1 | CPED1 | 1.24E-08 |
| pannexin 2 | PANX2 | 4.61E-08 |
| aldo-keto reductase family 1 member C2 | AKR1C2 | 3.41E-07 |
| dual specificity phosphatase 7 | DUSP7 | 9.34E-07 |
| Kruppel like factor 15 | KLF15 | 9.65E-07 |
| acyl-CoA dehydrogenase short/branched chain | ACADSB | 2.71E-06 |
| lectin, mannose binding 1 | LMAN1 | 3.11E-06 |
| PPARG coactivator 1 alpha | PPARGC1A | 5.70E-06 |
| aldo-keto reductase family 1 member B15 | AKR1B15 | 1.49E-05 |
| solute carrier family 22 member 31 | SLC22A31 | 2.67E-05 |
| TBC1 domain family member 4 | TBC1D4 | 3.26E-05 |
| C2 calcium dependent domain containing 2 | C2CD2 | 3.59E-05 |
| cingulin like 1 | CGNL1 | 5.32E-05 |
| solute carrier family 6 member 14 | SLC6A14 | 5.32E-05 |
| RNA, variant U1 small nuclear 31 | RNVU1-31 | 6.62E-05 |
| lysozyme | LYZ | 6.88E-05 |
| GDP-mannose 4,6-dehydratase | GMDS | 9.69E-05 |
| albumin | ALB | 1.07E-04 |
| FAST kinase domains 5 | FASTKD5 | 1.47E-04 |
| ferritin light chain | FTL | 1.48E-04 |
| TUB bipartite transcription factor | TUB | 1.57E-04 |
| suppressor APC domain containing 2 | SAPCD2 | 1.91E-04 |
| collagen type XXVI alpha 1 chain | COL26A1 | 2.00E-04 |

**TABLE 3.** Promoter analysis of downregulated genes in shFOXA1/2 knockdown HEPG2 cells.

| *Gene Symbol* | *Gene Name* | *Promoter-Liver GRN* | *Promoter-Other GRN* |
| --- | --- | --- | --- |
| AKR1C1 | aldo-keto reductase family 1 member C1 | FoxA1, FoxA2 | MRF-2, p53 |
| ALB | albumin | FoxA1, FoxA2, HNF4α, HNF4α | C/EBPalpha, STAT1, STAT1alpha, STAT1beta, STAT2, STAT3, STAT4, STAT5A, STAT5B, TBP |
| APOA4 | apolipoprotein A4 | FoxA1, FoxA2, HNF4α | COUP, COUP-TF, COUP-TF1, PPAR-gamma1, PPAR-gamma2 |
| APOC3 | apolipoprotein C3 | FoxA1, FoxA2, HNF1α, HNF4α | COUP, COUP-TF, COUP-TF1, PPAR-gamma1, PPAR-gamma2 |
| APOM | apolipoprotein M | FoxA1, FoxA2, HNF1α, HNF4α | COUP, COUP-TF, COUP-TF1, PPAR-gamma1, PPAR-gamma2, STAT1 |
| CD24 | CD24 molecule |  | c-Myc, FOXC1, Max1, Nkx3-1, Nkx3-1 v1, Nkx3-1 v2, Nkx3-1 v3, POU3F2, SRY |
| CYP19A1 | cytochrome P450 family 19 subfamily A member 1 | FoxA1, FoxA2, HNF4α | PPAR-gamma1, PPAR-gamma2 |
| CYP1A1 | cytochrome P450 family 1 subfamily A member 1 | FoxA1, FoxA2, HNF4α | Bach2, C/EBPbeta, GR, GR-alpha, MZF-1, p53, SREBP-1a, SREBP-1b, SREBP-1c |
| CYP24A1 | cytochrome P450 family 24 subfamily A member 1 | FoxA1, FoxA2, HNF1α, HNF4α | AML1a, AP-1, AREB6, ATF-2, c-Jun, C/EBPalpha, CREB, deltaCREB, FOXD3, YY1 |
| CYP4F2 | cytochrome P450 family 4 subfamily F member 2 | HNF4α | FOXO4, GATA-2, HOXA5, HSF1 (long), p53, Pax-2, Pax-2a, RORalpha1, STAT3, YY1 |
| DLK1 | delta like non-canonical Notch ligand 1 | FoxA1, FoxA2, HNF1α, HNF4α | Evi-1, GATA-2, PPAR-gamma1, PPAR-gamma2 |
| FASTKD5 | FAST kinase domains 5 | FoxA1, FoxA2, HNF4α | c-Myb, CP2, GATA-2, Ik-2, MZF-1, NF-kappaB, NF-kappaB1, Pax-2, Pax-2a, Zic3 |
| FGR1 | fibroblast growth factor receptor 1 | FoxA1, FoxA2, HNF1α, HNF4α | CREB, deltaCREB |
| IRS2 | insulin receptor substrate 2 | FoxA1, FoxA2, HNF4α | AP-1, ATF-2, c-Jun, C/EBPalpha, E2F, NRSF form 1, NRSF form 2 |
| KRT7 | keratin 7 | FoxA1,<br>FoxA2,<br>HNF4α | AP-1, c-Fos, c-Jun, p53, PPAR-gamma1, PPAR-gamma2 |
| NRARP | NOTCH-regulated ankyrin<br>repeat protein | FoxA1,<br>FoxA2,<br>HNF4α | Oct-B1, oct-B2, oct-B3, POU2F1,<br>POU2F1a, POU2F2, POU2F2 (Oct-2.1), POU2F2B, PPAR-gamma2 |
| PPARGC1A | PPARG coactivator 1 alpha | FoxA1,<br>FoxA2,<br>HNF4α | aMEF-2, CREB, deltaCREB, MEF-2A |
| TGFBR3 | transforming growth factor<br>beta receptor 3 | FoxA1,<br>FoxA2,<br>HNF1α,<br>HNF4α | c-Myc, GCNF-2, Max1, MyoD, Pax-2,<br>Pax-2a, Pax-2b, Sox9, Sp1 |
| TWNK | twinkle mtDNA helicase | FoxA1,<br>FoxA2,<br>HNF4α | |
| UCA1 | urothelial cancer assoc. 1 | FoxA1 |  |

**Figure 3.**
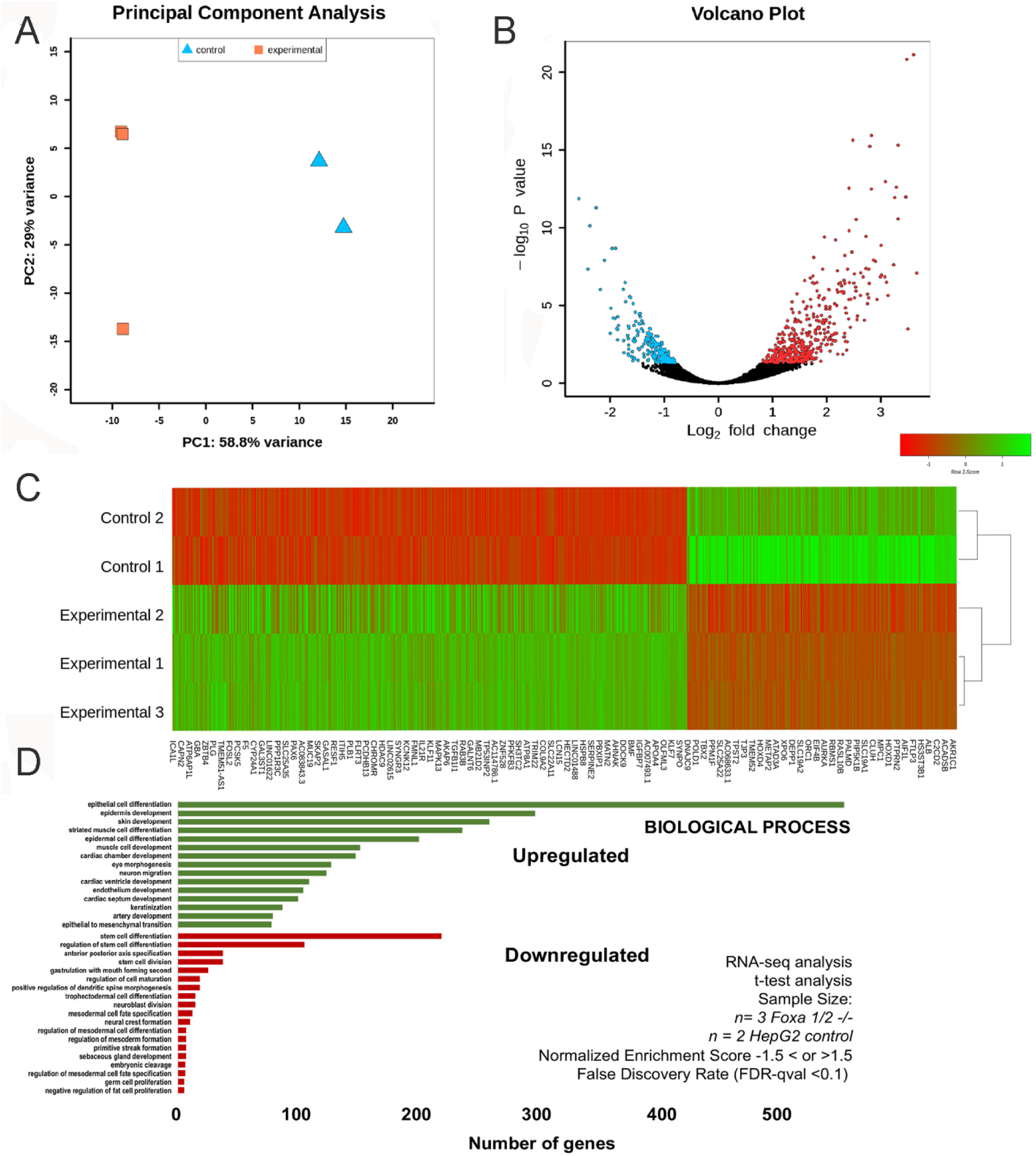
RNA-seq analysis of control (HepG2) and experimental (shFOXA1/2 knockdown) cells. A) Principal component analysis with experimental samples (n = 3) (orange), and two control samples (n = 2) (blue). Experimental group (shFOXA1/2) and control group are shown to cluster apart on the PC1 axis. B) Volcano plot displaying differential expression analysis between the control and experimental group. The adjusted p-value from the “res” function in DESeq2 was used as the p-value for this plot. Each circle in the plot represents a gene. The genes are colored if their adjusted p-value is below 0.05, which was used to indicate significant differential expression. The genes in green represent upregulation and the genes in red represent down regulation based on the log2-fold change. C) Heat map of differentially expressed genes for the five samples. Green represents relatively higher gene expression, red represents relatively lower gene expression, compared to other samples. Genes with less significant expression level differences (adjusted p-value > 0.05) were categorized as not differentially expressed and therefore not included. 448 upregulating and 235 downregulating genes are grouped together and listed by increasing adjusted P value for each group. Every seventh gene in the list is labeled. Hierarchical clustering was performed using the “hclust” function in R. D) Plot of biological process gene sets selected for relationship to stem cells and differentiation. Bioinformatics analysis of RNA seq data in which number of genes within each significant gene set is shown after removing genes not in the expression dataset. Red indicates a negative normalized enrichment score value or downregulation, green indicates a positive NES value or upregulation. Gene sets are from version 7.0 of the Gene Ontology (GO) gene sets from the Broad Institute. Number of genes within group shown after calculation, t-test comparison between groups (P < 0.05), for n = 2 control, n = 3 shFOXA1/2 cells. Criteria was NES -1.5 > or < 1.5, and false discovery rate (FDR)-qval < 0.1).

### Global transcriptome analysis of shFOXA1/2 cell phenotype demonstrates downregulation of metabolic genes associated with glycolysis and carbohydrate metabolism

We hypothesized that shFOXA1/2 cells exhibit global effects on cellular metabolism ^53^. We analyzed RNA-seq data to obtain a normalized enrichment score (NES) with a t-test, and obtained lists of up and down genes, from Gene Ontology (GO) and KEGG pathways **(Fig. 4A, Supp. Files 4-5**). Surprisingly, we observed hundreds of genes associated with nitrogen metabolism, oxygen reduction processes, and precursor metabolites **(Fig. 4A).** To confirm this, we analyzed gene expression of key enzymes involved in glucose metabolism. The data demonstrated significant downregulation of HK2, LDHA, PFKL, and PKM, with decrease of ALDOA approaching significance **(Fig. 4B-C)** in shFOXA1/2 knockdown cells. Interestingly, in FOXA2 depleted cells, we found that there was no change in ALDOA, LDHA, PFKL, and PKM gene expression in FOXA2 depleted cells, indicating that FOXA1 may have compensate. To further elicit metabolic changes, we mapped gene expression data for glycolysis and gluconeogenesis **(Fig. 4D)**, citric acid cycle **(Fig. 4E)**, glycogen formation and breakdown **(Fig. 4F)**, and pentose phosphate pathway **(Fig. 4G-H)**. Overall, for glycolysis and gluconeogenesis we observed 31 downregulated (9 significant, 22 trending) and 22 upregulated (4 significant, 18 trending) (**Table 4**). For example, downregulated (trending) genes included hexokinase (HK2) and phosphofructokinase-1 (PFKL), known to be rate-limiting enzymes, as well as triose phosphate isomerase (TPI1), and glyceraldehyde phosphate dehydrogenase (GAPDH). Downregulated (significant) genes included phosphohexose isomerase (GPI). These data suggest a downregulation of glycolysis and buildup of intermediates, like Fructose 6-Phosphate. Next, we analyzed genes of the Tricarboxylic (TCA) or citric acid cycle **(Fig. 4E)**, and overall, we found 26 genes were downregulated (9 significant, 17 trending), and 4 genes were upregulated (1 significant, 3 trending) (**Table 5**). Significantly downregulated genes included 2/3 isoforms of pyruvate dehydrogenases (PDHA1, PDHB), 1/2 isoforms of aconitase (ACO2), 2/5 isoforms of Isocitrate Dehydrogenases (IDH1, IDH3B), 1/3 of isoforms of Succinyl-CoA Synthetase (SUCLG2), 1/4 isoforms of Succinate Dehydrogenase (SDHB), and Fumarase (PH). These data suggest a near complete downregulation of the TCA cycle. For glycogenolysis **(Fig. 4F)**, we found 13 genes downregulated (1 significant, 12 trending), and 2 upregulated (1 significant, 1 trending), (**Table 6**). For glycogen synthesis ((**Fig. 4F)**, we found 10 genes downregulated (4 significant, 6 trending), and 5 upregulated (2 significant, 3 trending) (**Table 6**). For Non-oxidative and Oxidative Pentose Phosphate Pathways, we observed 15 downregulated (6 significant, 9 trending), and 9 upregulated (1 significant, 8 trending) **(Fig. 4G-H, Table 7)**. GLUT1 transporter has been shown to uptake glucose in HepG2 cells ^54^ but was not differentially regulated in our system (data not shown). Based on the widespread changes in global gene expression associated with glucose metabolism in the shFOXA1/2 cells, we measured extracellular acidification rate (ECAR) which is an indicator of glycolysis using Seahorse Analyzer in control and FOXA1/2 depleted HepG2 in the presence of selected glycolytic inhibitors, oligomycin and 2-deoxyglucose **(Fig. 4I)**. We found that glycolysis (n = 3, p = 0.042) and glycolytic capacity (n = 3, p = 0.024) were significantly decreased in the shFOXA1/2 cells **(Fig. 4J-K).** Overall, we demonstrated several downregulated genes in key pathways indicating that glycolytic and TCA cycle intermediates were likely pooling, and that carbons from glucose were not flowing through the TCA cycle.

**Figure 4.**
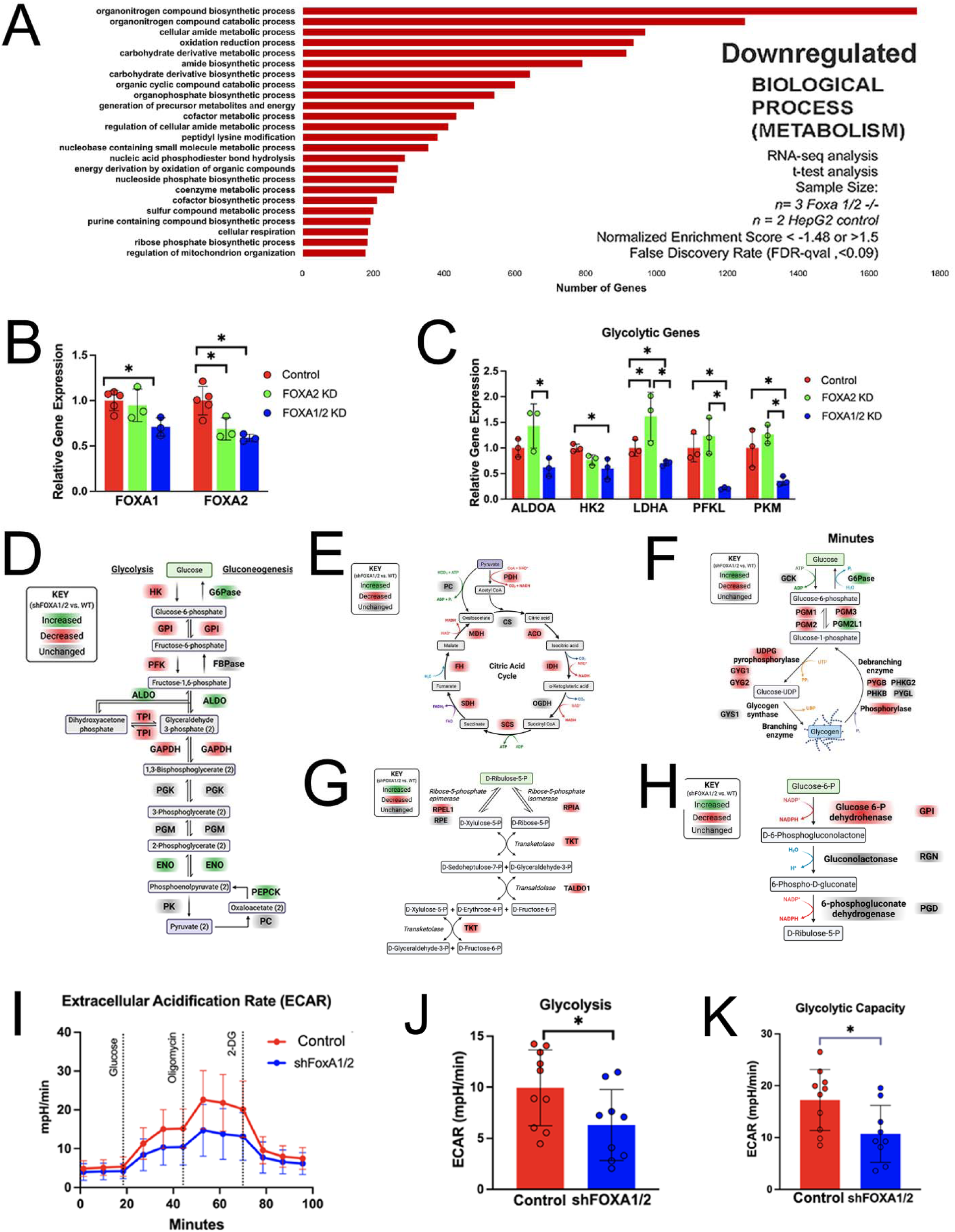
RNA-seq analysis metabolism of control (HepG2) and experimental (shFOXA1/2 knockdown) cells. A) Plot of biological process gene sets selected for relationship to metabolism. Bioinformatics analysis of RNA seq data in which number of genes within each significant gene set is shown after removing genes not in the expression dataset. Red indicates a negative normalized enrichment score value or downregulation, green indicates a positive NES value or upregulation. Gene sets are from version 7.0 of the Gene Ontology (GO) gene sets from the Broad Institute. Number of genes within group shown after calculation, t-test comparison between groups (P < 0.05), for n = 2 control, n = 3 shFOXA1/2 cells. Criteria was NES -1.48 > or < 1.5, and false discovery rate (FDR)-qval < 0.09). B) Bar graph of gene expression (qRT-PCR) of FOXA1 and FOXA2 in control (scrambled shRNA control) and shFOXA1/2 knockdown cells. N = 3 for all conditions. Plotted is mean ± SD. Significance (*) defined as P ≤ 0.05. C) Bar graph of gene expression (qRT-PCR) of glycolysis associated genes in control (scrambled shRNA control) and shFOXA1/2 knockdown cells. N = 3 for all conditions. Plotted is mean ± SD. Significance (*) defined as P ≤ 0.05. D) Schematic map of glycolysis and gluconeogenesis demonstrating downregulated (red) and upregulated (green) pathways. Downregulated genes include downtrending genes (p < 0.2) and downregulated (p < 0.05). Uptrending genes (p < 0.20) and upregulated genes (p < 0.05). E) Same as D except tricarboxylic cycle (cycle acid cycle) F) Same as D except glycogen synthesis and glycogenolysis G) Same as D except non-oxidative pentose phosphate pathway H) Same as D except oxidative pentose phosphate pathway I) Functional analysis of glycolysis in control (scrambled shRNA control) and shFOXA1/2 knockdown cells. Plot of extracellular acidification rate (ECAR) versus time using Seahorse XL. Data demonstrates effects of addition of glucose, oligomycin (ATP synthetase inhibitor), and 2-DG (2-Deoxy-d-glucose, glycolysis inhibitor). J) Bar graph measuring glycolysis rate (ECAR) in control (scrambled shRNA control) and shFOXA1/2 knockdown cells. N = 3 for all conditions. Plotted is mean ± SD. Significance (*) defined as P ≤ 0.05. K) Bar graph measuring glycolytic capacity (ECAR) in control (scrambled shRNA control) and shFOXA1/2 knockdown cells. * indicates significance (p <0.05). N = 3 for all conditions. Plotted is mean ± SD. Significance (*) defined as P ≤ 0.05.

**TABLE 4.**
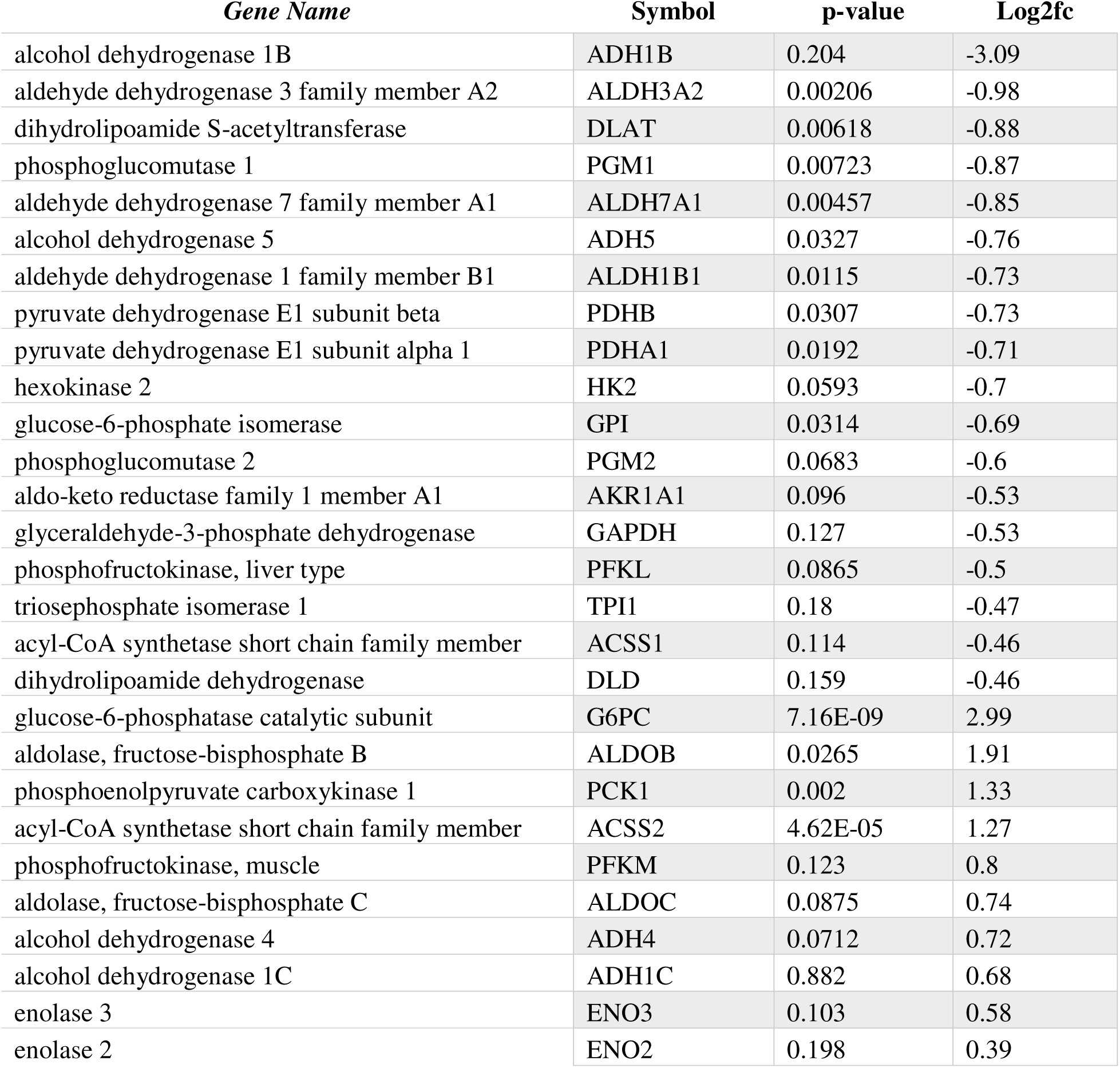
Glycolysis and gluconeogenesis pathway genes in shFOXA1/2 knockdown HEPG2 cells.

| *Gene Name* | *Symbol* | *p-value* | *Log2fc* |
| --- | --- | --- | --- |
| alcohol dehydrogenase 1B | ADH1B | 0.204 | -3.09 |
| aldehyde dehydrogenase 3 family member A2 | ALDH3A2 | 0.00206 | -0.98 |
| dihydrolipoamide S-acetyltransferase | DLAT | 0.00618 | -0.88 |
| phosphoglucomutase 1 | PGM1 | 0.00723 | -0.87 |
| aldehyde dehydrogenase 7 family member A1 | ALDH7A1 | 0.00457 | -0.85 |
| alcohol dehydrogenase 5 | ADH5 | 0.0327 | -0.76 |
| aldehyde dehydrogenase 1 family member B1 | ALDH1B1 | 0.0115 | -0.73 |
| pyruvate dehydrogenase E1 subunit beta | PDHB | 0.0307 | -0.73 |
| pyruvate dehydrogenase E1 subunit alpha 1 | PDHA1 | 0.0192 | -0.71 |
| hexokinase 2 | HK2 | 0.0593 | -0.7 |
| glucose-6-phosphate isomerase | GPI | 0.0314 | -0.69 |
| phosphoglucomutase 2 | PGM2 | 0.0683 | -0.6 |
| aldo-keto reductase family 1 member A1 | AKR1A1 | 0.096 | -0.53 |
| glyceraldehyde-3-phosphate dehydrogenase | GAPDH | 0.127 | -0.53 |
| phosphofructokinase, liver type | PFKL | 0.0865 | -0.5 |
| triosephosphate isomerase 1 | TPI1 | 0.18 | -0.47 |
| acyl-CoA synthetase short chain family member | ACSS1 | 0.114 | -0.46 |
| dihydrolipoamide dehydrogenase | DLD | 0.159 | -0.46 |
| glucose-6-phosphatase catalytic subunit | G6PC | 7.16E-09 | 2.99 |
| aldolase, fructose-bisphosphate B | ALDOB | 0.0265 | 1.91 |
| phosphoenolpyruvate carboxykinase 1 | PCK1 | 0.002 | 1.33 |
| acyl-CoA synthetase short chain family member | ACSS2 | 4.62E-05 | 1.27 |
| phosphofructokinase, muscle | PFKM | 0.123 | 0.8 |
| aldolase, fructose-bisphosphate C | ALDOC | 0.0875 | 0.74 |
| alcohol dehydrogenase 4 | ADH4 | 0.0712 | 0.72 |
| alcohol dehydrogenase 1C | ADH1C | 0.882 | 0.68 |
| enolase 3 | ENO3 | 0.103 | 0.58 |
| enolase 2 | ENO2 | 0.198 | 0.39 |

**TABLE 5.** Citrate Cycle pathway genes in shFOXA1/2 knockdown HEPG2 cells.

| *Gene Name* | **Symbol** | **p-value** | **Log2fc** |
| --- | --- | --- | --- |
| succinate-CoA ligase GDP-forming subunit b | SUCLG2 | 0.000614 | -1.16 |
| dihydrolipoamide S-acetyltransferase | DLAT | 0.00618 | -0.88 |
| succinate dehydrogenase complex iron sulfur | SDHB | 0.0111 | -0.87 |
| fumarate hydratase | FH | 0.013 | -0.87 |
| isocitrate dehydrogenase | IDH3B | 0.00467 | -0.85 |
| pyruvate dehydrogenase E1 subunit beta | PDHB | 0.0307 | -0.73 |
| aconitase 2 | ACO2 | 0.0174 | -0.72 |
| pyruvate dehydrogenase E1 subunit alpha 1 | PDHA1 | 0.0192 | -0.71 |
| isocitrate dehydrogenase | IDH1 | 0.0354 | -0.68 |
| isocitrate dehydrogenase | IDH3A | 0.0734 | -0.66 |
| succinate dehydrogenase complex subunit D | SDHD | 0.0948 | -0.63 |
| succinate-CoA ligase GDP/ADP-forming subunit | SUCLG1 | 0.0748 | -0.62 |
| malate dehydrogenase 2 | MDH2 | 0.0534 | -0.6 |
| malate dehydrogenase 1 | MDH1 | 0.0648 | -0.6 |
| succinate-CoA ligase ADP-forming subunit | SUCLA2 | 0.0922 | -0.53 |
| dihydrolipoamide S-succinyltransferase | DLST | 0.116 | -0.5 |
| succinate dehydrogenase complex flavoprotein | SDHA | 0.153 | -0.5 |
| dihydrolipoamide dehydrogenase | DLD | 0.159 | -0.46 |
| succinate dehydrogenase complex subunit C | SDHC | 0.144 | -0.42 |
| phosphoenolpyruvate carboxykinase 1 | PCK1 | 0.002 | 1.33 |

**TABLE 6.** Glycogenolysis and glycogen formation genes in shFOXA1/2 knockdown HEPG2 cells.

| *Gene Name* | **Symbol** | **p-value** | **Log2fc** |
| --- | --- | --- | --- |
| glycogenin 2 | GYG2 | 5.12E-05 | -1.29 |
| phosphoglucomutase 1 | PGM1 | 0.00723 | -0.87 |
| phosphoglucomutase 2=3 | PGM3 | 0.026 | -0.78 |
| ubiquitin A-52 residue ribosomal protein | UBA52 | 0.0246 | -0.76 |
| hexokinase 2 | HK2 | 0.0593 | -0.7 |
| ribosomal protein S27a | RPS27A | 0.0434 | -0.68 |
| phosphoglucomutase 2 | PGM2 | 0.0683 | -0.6 |
| calmodulin 1 | CALM1 | 0.0917 | -0.58 |
| glycogenin 1 | GYG1 | 0.158 | -0.56 |
| glycogen phosphorylase B | PYGB | 0.0975 | -0.48 |
| ubiquitin C | UBC | 0.179 | -0.4 |
| glucosidase alpha, acid | GAA | 0.216 | -0.36 |
| glucose-6-phosphatase catalytic subunit | G6PC | 7.16E-09 | 2.99 |
| NHL repeat containing E3 ubiquitin | NHLRC1 | 0.144 | 1.88 |
| phosphoglucomutase 2 like 1 | PGM2L1 | 0.000266 | 1.5 |
| protein phosphatase 1 regulatory subunit | PPP1R3C | 0.000754 | 1.46 |

**TABLE 7.** Pentose Phosphate Pathway genes in shFOXA1/2 knockdown HEPG2 cells.

| *Gene Name* | **Symbol** | **p-value** | **Log2fc** |
| --- | --- | --- | --- |
| ribose 5-phosphate isomerase A | RPIA | 0.00285 | -1.28 |
| transaldolase 1 | TALDO1 | 0.000726 | -1.1 |
| ribulose-5-phosphate-3-epimerase like 1 | RPEL1 | 0.203 | -1.09 |
| phosphoribosyl pyrophosphate synthetase 1 | PRPS1 | 0.00612 | -0.92 |
| phosphoglucomutase 1 | PGM1 | 0.00723 | -0.87 |
| transketolase | TKT | 0.0197 | -0.76 |
| glucose-6-phosphate isomerase | GPI | 0.0314 | -0.69 |
| phosphoglucomutase 2 | PGM2 | 0.0683 | -0.6 |
| phosphofructokinase, liver type | PFKL | 0.0865 | -0.5 |
| phosphoribosyl pyrophosphate synthetase 2 | PRPS2 | 0.16 | -0.48 |
| aldolase, fructose-bisphosphate B | ALDOB | 0.0265 | 1.91 |
| phosphofructokinase, muscle | PFKM | 0.123 | 0.8 |
| aldolase, fructose-bisphosphate C | ALDOC | 0.0875 | 0.74 |

### Functional metabolic analysis of shFOXA1/2 liver cell line demonstrates defects in mitochondrial metabolism

Given the widespread metabolic changes in shFOXA1/2 cells, we used the same upregulated and downregulated gene sets for Gene Ontology (GO) and KEGG pathways for genes associated with mitochondrial metabolism **(Fig. 5A)**. Amazingly, we identified approximately 1500 mitochondria-related genes, including 1000 genes associated with mitochondrial envelope, over 450 genes associated with mitochondrial matrix, and over 200 genes associated with mitochondrial protein complex **(Fig. 5A)**. We mapped trending (p < 0.20) and significant (p < 0.05) downregulated and upregulated genes onto a schematic that demonstrated the mitochondrial electron transport chain. Overall, we observed 53 downregulated (15 significant, 38 trending) and 8 upregulated (2 significant, 6 trending) genes of interest, including subtypes or isoforms **(Fig. 5B, Table 8**). For Complexes I-V in the electron transport chain, we observed several downregulated genes. For example, for Complex V (ATP synthesis), we observed downregulation of ATP5F1A, ATP5F1D, ATP5F1E, ATP5MC1, ATP5MC3, and ATP5PB, (significant) and ATP5F1C and ATP5PF (trending) **(Fig. 5B, Table 8)**. We concluded that there was a lack of energy production via mitochondrial metabolism. Since mitochondrial metabolism is globally regulated by peroxisome proliferator-activated receptor γ (PPARγ) co-activator (PGC1α) ^55–56^ and is associated with insulin resistance and non-alcoholic steatohepatitis, we examined its expression levels. We found that PGC1α was downregulated in shFOXA1/2 knockdown cells (**Table 2**). However, genes associated with PGC1α, including CREB, FOXO1, SREBP, and were not downregulated (data not shown). To further assess metabolic deficits, we examined the urea cycle (nitrogen and ammonia metabolism), which generates carbon skeletons for energy generation. We suspected that shFOXA1/2 cells may be using aspects of the urea cycle as a carbon source. We observed a trending increase in urea production **(Fig. 2I)** and a downregulation of 3 urea cycle genes (1 trending, 2 significant, **Table 9)**. Based on our = increased fat droplets **(Fig. 2H)**, we examined genes associated with lipid ketogenesis and lipid metabolism (**Tables 10-11**). In ketone pathways in shFOXA1/2 cells, and we found 9 genes upregulated (1 significant, 2 trending), and 5 genes downregulated (1 significant, 2 trending) (**Table 10**). We examined lipid metabolism and found 13 genes upregulated (2 significant, 1 trending) and 27 downregulated (11 significant, 4 trending) (**Table 11**). This data could explain the buildup of fat droplets we observed in **(Fig. 2G)**. To identify potential sources of carbon, we examined nucleo-cytosolic acetyl-CoA synthetase enzyme (ACSS2) expression which has been associated with tumor cells that upregulate acetate uptake ^57^ and we observed it was significantly upregulated in the shFOXA1/2 population, suggesting the potential of acetate uptake in these cells. Overall, the data suggests mechanisms for changes in lipid metabolism and acetate uptake/metabolism suggest that lipids, and extracellular or intracellular acetate could be involved as sources of carbon for oxidation, energy generation, or anabolic reactions. We therefore hypothesized that there are functional changes. We measured oxygen consumption rate (OCR), which is an indicator of mitochondrial respiration, using Seahorse Analyzer in control and FOXA1/2 depleted HepG2 with glycolytic inhibitors oligomycin, FCCP, or Rotenone **(Fig. 5C)**. We found that basal respiration (n = 3, p = 0.042), ATP-linked respiration (n = 3, p = 0.042), and proton leak (n = 3, p = 0.042) were significantly decreased in the shFOXA1/2 cells **(Fig. 5D-F)**. There were no differences in maximal respiration (data not shown). Collectively, these data suggest broad dysfunction in mitochondrial respiration in the shFOXA1/2 cells, and that FOXA1 can potentially compensate for FOXA2 for glycolytic enzymes.

**Figure 5.**
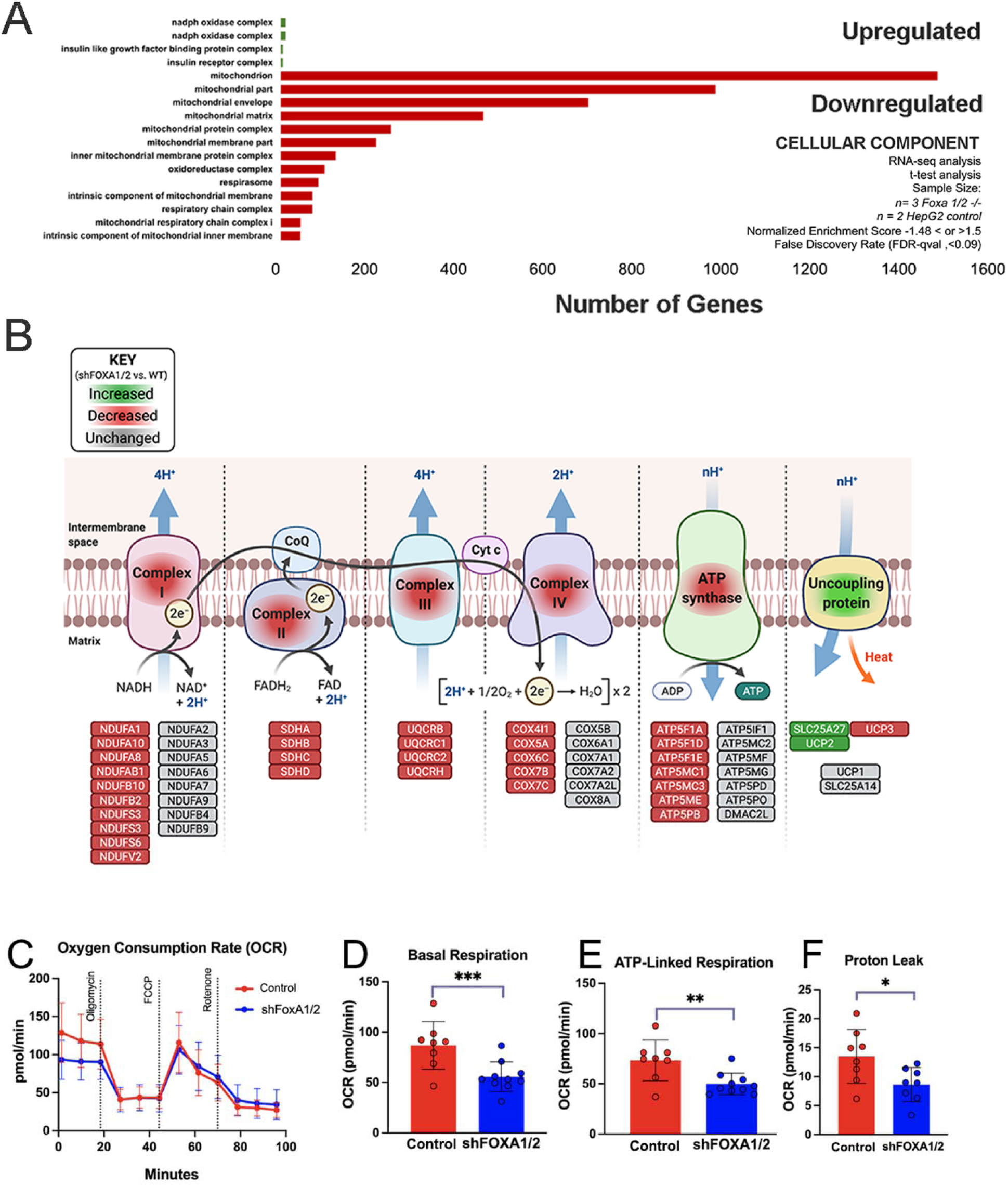
Functional metabolic analysis of the shFOXA1/2 liver cell line. A) Plot of cellular component gene sets selected for relationship to metabolism. Bioinformatics analysis of RNA seq data in which number of genes within each significant gene set is shown after removing genes not in the expression dataset. Red indicates a negative normalized enrichment score value or downregulation, green indicates a positive NES value or upregulation. Gene sets are from version 7.0 of the Gene Ontology (GO) gene sets from the Broad Institute. Number of genes within group shown after calculation, t-test comparison between groups (P < 0.05), for n = 2 control, n = 3 shFOXA1/2 cells. Criteria was NES -1.48 > or < 1.5, and false discovery rate (FDR)-qval < 0.09). B) Schematic map of genes associated with electron transport chain demonstrating downregulated (red) and upregulated (green) pathways. Downregulated genes include downtrending genes (p < 0.2) and downregulated (p < 0.05). Uptrending genes (p < 0.20) and upregulated genes (p < 0.05). Right-Map of citric acid cycle demonstrating upregulated and downregulated pathways. Same scheme for upregulation and downregulation is shown. C) Functional analysis of glycolysis in control (scrambled shRNA control) and shFOXA1/2 knockdown cells. Plot of oxygen consumption rate (OCR) versus time using Seahorse XL. Data demonstrates effects of addition of glucose, oligomycin (ATP synthetase inhibitor), and 2-DG (2-Deoxy-d-glucose, glycolysis inhibitor). D) Bar graph measuring basal respiration (OCR) in control (scrambled shRNA control) and shFOXA1/2 knockdown cells. N = 3 for all conditions. Plotted is mean ± SD. Significance (*) defined as P ≤ 0.05. E) Bar graph measuring Proton leak in control (scrambled shRNA control) and shFOXA1/2 knockdown cells. N = 3 for all conditions. Plotted is mean ± SD. Significance (*) defined as P ≤ 0.05. F) Bar graph measuring ATP-linked respiration in control (scrambled shRNA control) and shFOXA1/2 knockdown cells. N = 3 for all conditions. Plotted is mean ± SD. Significance (*) defined as P ≤ 0.05.

**TABLE 8.**
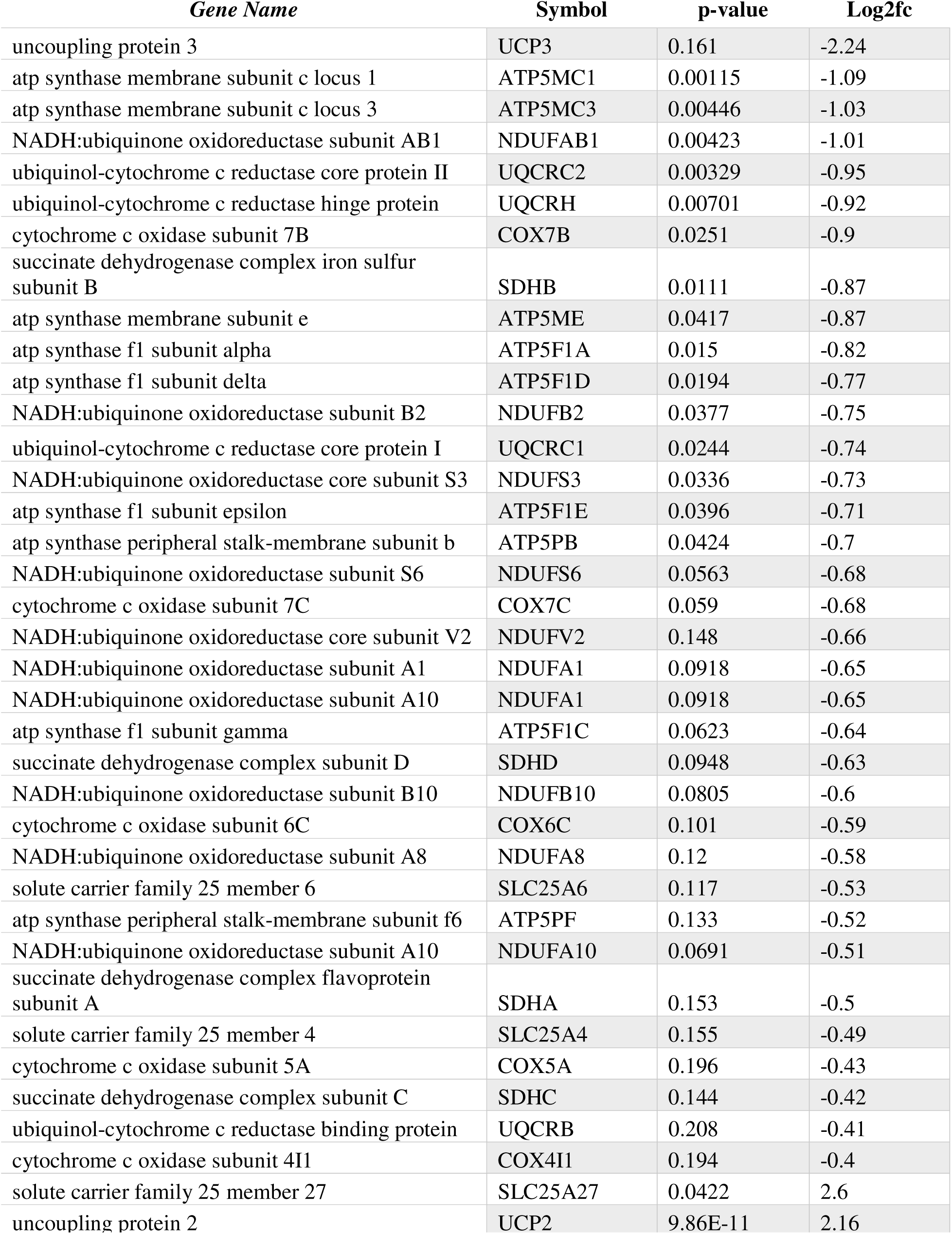
Electron Transport Chain genes in shFOXA1/2 knockdown HEPG2 cells.

| *Gene Name* | *Symbol* | *p-value* | *Log2fc* |
| --- | --- | --- | --- |
| uncoupling protein 3 | UCP3 | 0.161 | -2.24 |
| atp synthase membrane subunit c locus 1 | ATP5MC1 | 0.00115 | -1.09 |
| atp synthase membrane subunit c locus 3 | ATP5MC3 | 0.00446 | -1.03 |
| NADH:ubiquinone oxidoreductase subunit AB1 | NDUFAB1 | 0.00423 | -1.01 |
| ubiquinol-cytochrome c reductase core protein II | UQCRC2 | 0.00329 | -0.95 |
| ubiquinol-cytochrome c reductase hinge protein | UQCRH | 0.00701 | -0.92 |
| cytochrome c oxidase subunit 7B | COX7B | 0.0251 | -0.9 |
| succinate dehydrogenase complex iron sulfur subunit B | SDHB | 0.0111 | -0.87 |
| atp synthase membrane subunit e | ATP5ME | 0.0417 | -0.87 |
| atp synthase f1 subunit alpha | ATP5F1A | 0.015 | -0.82 |
| atp synthase f1 subunit delta | ATP5F1D | 0.0194 | -0.77 |
| NADH:ubiquinone oxidoreductase subunit B2 | NDUFB2 | 0.0377 | -0.75 |
| ubiquinol-cytochrome c reductase core protein I | UQCRC1 | 0.0244 | -0.74 |
| NADH:ubiquinone oxidoreductase core subunit S3 | NDUFS3 | 0.0336 | -0.73 |
| atp synthase f1 subunit epsilon | ATP5F1E | 0.0396 | -0.71 |
| atp synthase peripheral stalk-membrane subunit b | ATP5PB | 0.0424 | -0.7 |
| NADH:ubiquinone oxidoreductase subunit S6 | NDUFS6 | 0.0563 | -0.68 |
| cytochrome c oxidase subunit 7C | COX7C | 0.059 | -0.68 |
| NADH:ubiquinone oxidoreductase core subunit V2 | NDUFV2 | 0.148 | -0.66 |
| NADH:ubiquinone oxidoreductase subunit A1 | NDUFA1 | 0.0918 | -0.65 |
| NADH:ubiquinone oxidoreductase subunit A10 | NDUFA1 | 0.0918 | -0.65 |
| atp synthase f1 subunit gamma | ATP5F1C | 0.0623 | -0.64 |
| succinate dehydrogenase complex subunit D | SDHD | 0.0948 | -0.63 |
| NADH:ubiquinone oxidoreductase subunit B10 | NDUFB10 | 0.0805 | -0.6 |
| cytochrome c oxidase subunit 6C | COX6C | 0.101 | -0.59 |
| NADH:ubiquinone oxidoreductase subunit A8 | NDUFA8 | 0.12 | -0.58 |
| solute carrier family 25 member 6 | SLC25A6 | 0.117 | -0.53 |
| atp synthase peripheral stalk-membrane subunit f6 | ATP5PF | 0.133 | -0.52 |
| NADH:ubiquinone oxidoreductase subunit A10 | NDUFA10 | 0.0691 | -0.51 |
| succinate dehydrogenase complex flavoprotein subunit A | SDHA | 0.153 | -0.5 |
| solute carrier family 25 member 4 | SLC25A4 | 0.155 | -0.49 |
| cytochrome c oxidase subunit 5A | COX5A | 0.196 | -0.43 |
| succinate dehydrogenase complex subunit C | SDHC | 0.144 | -0.42 |
| ubiquinol-cytochrome c reductase binding protein | UQCRB | 0.208 | -0.41 |
| cytochrome c oxidase subunit 4I1 | COX4I1 | 0.194 | -0.4 |
| solute carrier family 25 member 27 | SLC25A27 | 0.0422 | 2.6 |
| uncoupling protein 2 | UCP2 | 9.86E-11 | 2.16 |

**TABLE 9.** Urea Cycle in shFOXA1/2 knockdown HEPG2 cells.

| *Gene Name* | **Symbol** | **p-value** | **Log2fc** |
| --- | --- | --- | --- |
| solute carrier family 25 member 1 | SLC25A15 | 0.0101 | -0.84 |
| argininosuccinate lyase | ASL | 0.0411 | -0.63 |
| ornithine carbamoyltransferase | OTC | 0.106 | -0.52 |

**TABLE 10.**
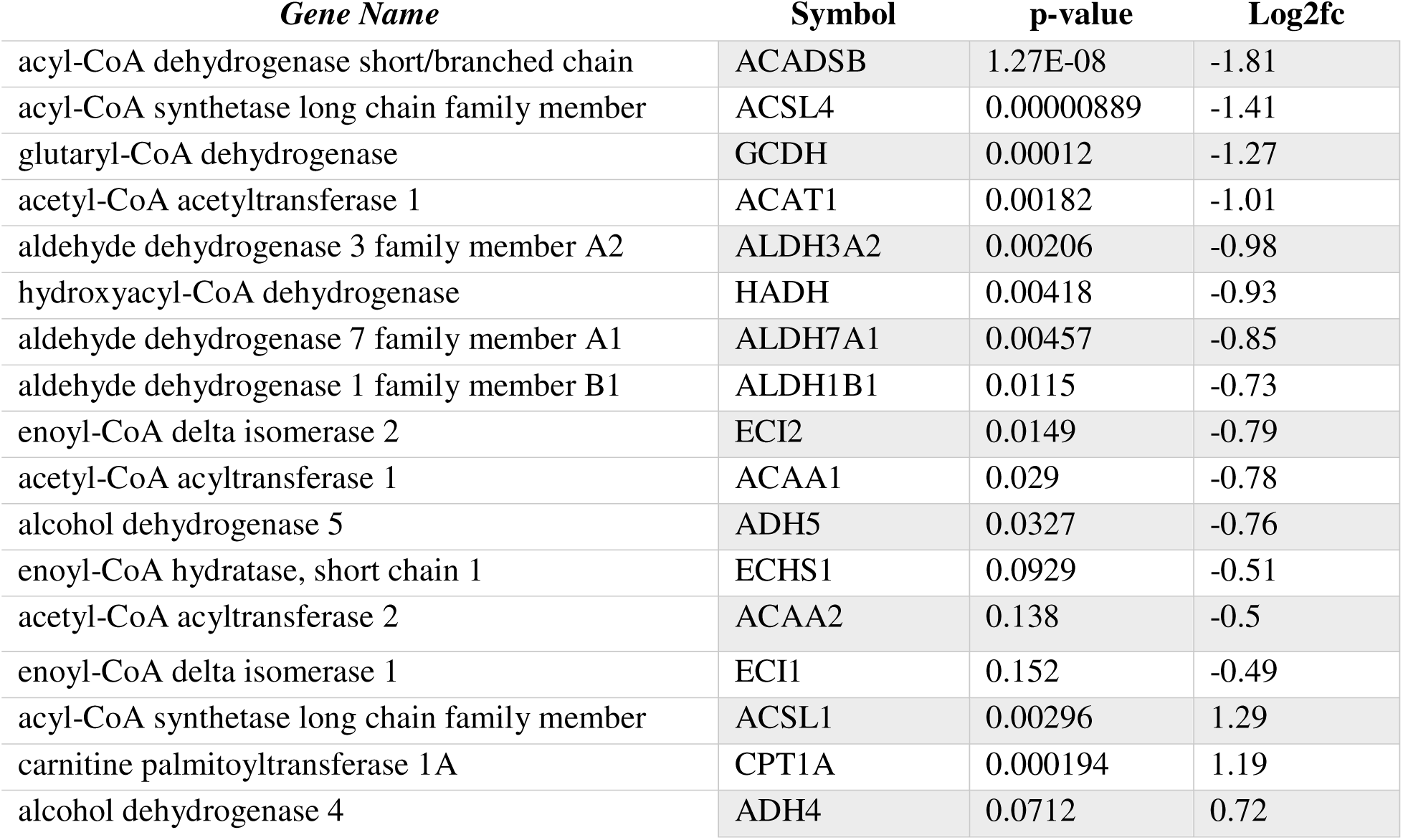
Ketogenesis in shFOXA1/2 knockdown HEPG2 cells.

**TABLE 11.** Lipogenesis in shFOXA1/2 knockdown HEPG2 cells.

| *Gene Name* | **Symbol** | **p-value** | **Log2fc** |
| --- | --- | --- | --- |
| acetyl-coa acetyltransferase 1 | ACAT1 | 0.00182 | -1.01 |
| acetoacetyl-coa synthetase | AACS | 0.0276 | -0.71 |
| solute carrier family 16 member 1 | SLC16A1 | 0.114 | -0.5 |
| uncoupling protein 2 | UCP2 | 9.86E-11 | 2.16 |
| 3-hydroxy-3-methylglutaryl-coa lyase | HMGCL | 0.0553 | 0.71 |
| 3-hydroxybutyrate dehydrogenase 1 | BDH1 | 0.112 | 0.61 |

### siFOXA1/2 and shFOXA1/2 knockdown arrests hepatic differentiation, downregulated hepatic GRN, and alters pluripotency factors in hPSC-derived liver differentiation

FOXA1/2 regulate the initiation of murine liver differentiation, but it is not known whether FOXA1/2 To answer this, we developed a human pluripotent stem cell (hPSC)-derived liver differentiation protocol that is focused on initiation of liver differentiation. hPSC (induced pluripotent stem cells (iPSC) and embryonic stem cells) at various passages were cultivated in pluripotent colonies **(Fig. 6A)** and subjected to our protocol for endoderm induction **(Fig. 6B-G)**. Under hypoxic conditions, we formed definitive endoderm that expressed FOXA2 and SOX17, with no ALB and CDX2 gene expression detected **(Fig. 6G)**. Our analysis of liver development indicated that for several weeks, hypoxic conditions remain ^58–60^. Therefore, we wished to differentiate hepatoblasts under continued hypoxic conditions. We first employed a traditional published protocol ^61^ and used it under hypoxic conditions, but observed poor morphology and gene expression. We compared a traditional liver differentiation protocol **(Fig. 6H)** to a new protocol (GF (-)) that we developed employing a nutrient rich, serum-free medium termed SFD, which has been used to maintain DE and enable spontaneous hepatic differentiation ^44^ over a total of 14 days **(Fig. 6H)**. Using this second protocol, our morphology, gene and protein expression studies demonstrate high levels of hepatic differentiation by day 14 in the GF (-) condition **(Fig. 6I-K)**.

**Figure 6.**
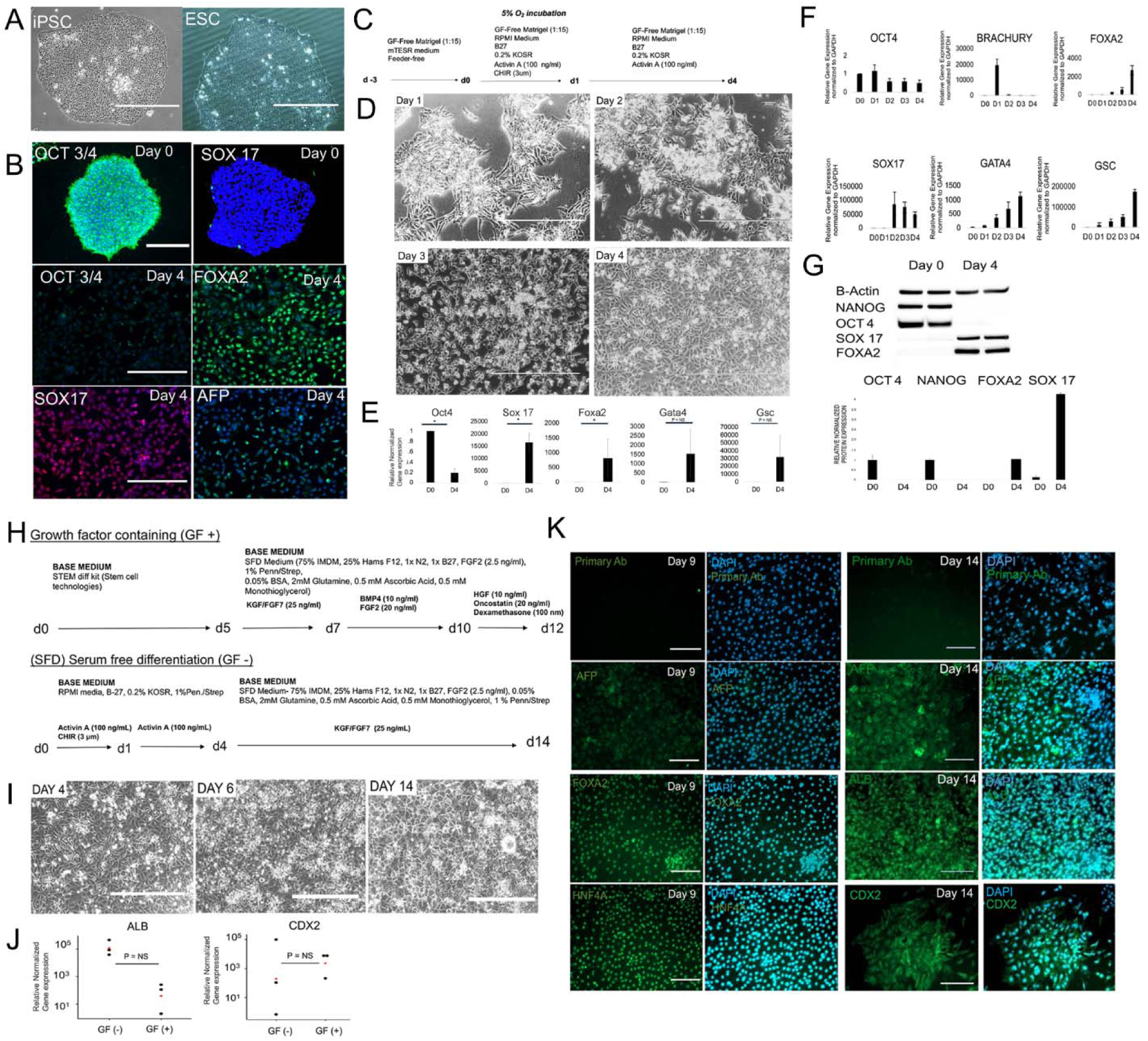
Spontaneously differentiated human stem cell-derived hepatic progenitors. A) Phase contrast images of Day 0 hPSC (iPSC and hESC). B) Schematic of protocol for definitive endoderm induction from hPSC under hypoxic conditions. C) Immunostaining of Day 0 (top, OCT4, SOX 17) and Day 4 definitive endoderm cells (OCT4, FOXA2, SOX17, AFP). D) Phase contrast microscopy during differentiation from Day 0 to Day 4 (endoderm), Bar = 400 µm. E) Bar graph of gene expression (qRT-PCR) on Day 0 and Day 4 during endoderm induction for OCT4, SOX17, FOXA2, GSC, GATA4, GSC. Plotted is mean ± SD. Significance (*) defined as P ≤ 0.05. F) Bar graph showing kinetics of gene expression (qRT-PCR) during endoderm induction for OCT4, BRACHYURY, FOXA2, SOX 17, GATA4, GSC. Plotted is mean ± SD for replicates at each time point (n = 1). G) Western blot of pluripotency (NANOG, OCT4) and definite endoderm (FOXA2, SOX 17) on Day 0 and Day 4 during definite endoderm induction. Quantitation of Western Blot below. H) Schematics of GF (+) and GF (-) liver differentiation protocols both under low oxygen conditions. GF (+) is adopted from Takebe et al. ^61^ and has the standard growth factors. The GF (-) protocol is a protocol based on spontaneous differentiation no additional factors added under than KGF which specifies gut tube from definitive endoderm. The protocol involves induction of endoderm by Day 5, and then switch of medium to SFD medium, followed by differentiation to gut tube by Day 7, and continues the addition of KGF with no additional growth factors from Day 7-Day 14, under hypoxic conditions. I) Phase contrast microscopy demonstrating morphology during differentiation from Day 4 (endoderm), Day 6 (gut tube endoderm), and Day 14 (definitive endoderm). Bar = 400 µm. J) Scatter plot of ALB gene expression (qRT-PCR) on Day 14 of differentiation, in the GF (-) condition compared to the GF (+) condition, on a log plot. N = 3 for both conditions. ALB, P = NS (P = 0.21). Plotted are individual values and mean. Significance (*) defined as P ≤ 0.05 (n = 3). K) Immunostaining of FOXA2, AFP, and HNF4A on Day 9. Bar = 200 µm.

Using this novel hepatoblast protocol, we performed siFOXA1/2 targeting on day 11 of hPSC hepatic differentiation. We identified downregulation of the liver gene expression, including FOXA1, FOXA2, AFP, and ALB **(Fig. 7A)**. We suspected that if differentiation was blocked, de-differentiation would occur. Surprisingly, we found a significant upregulation of OCT4, NANOG, and SOX2, suggesting that pluripotency factors were upregulated **(Fig. 7B)**. We also observed upregulation of PAX6 (neuroectoderm), no change in DE (SOX17), and HNF61B, and downregulation of CDX2, HNF4A, HHEX, PROX1 **(Fig. 7B-C)**. Further, pancreas endoderm, (PDX1) and lung endoderm (NKX2.1) were downregulated, while there was no change in hepatobiliary (SOX9), and biliary (CK19) **(Fig. 7D)**. Western blot confirmed downregulation of FOXA1/2, and ALB, and demonstrated an upregulation of NANOG and SOX2, as expected **(Fig. 7E-F)**. These data suggest an arrest of liver differentiation and an upregulation of pluripotency and neuroectodermal markers.

**Figure 7.**
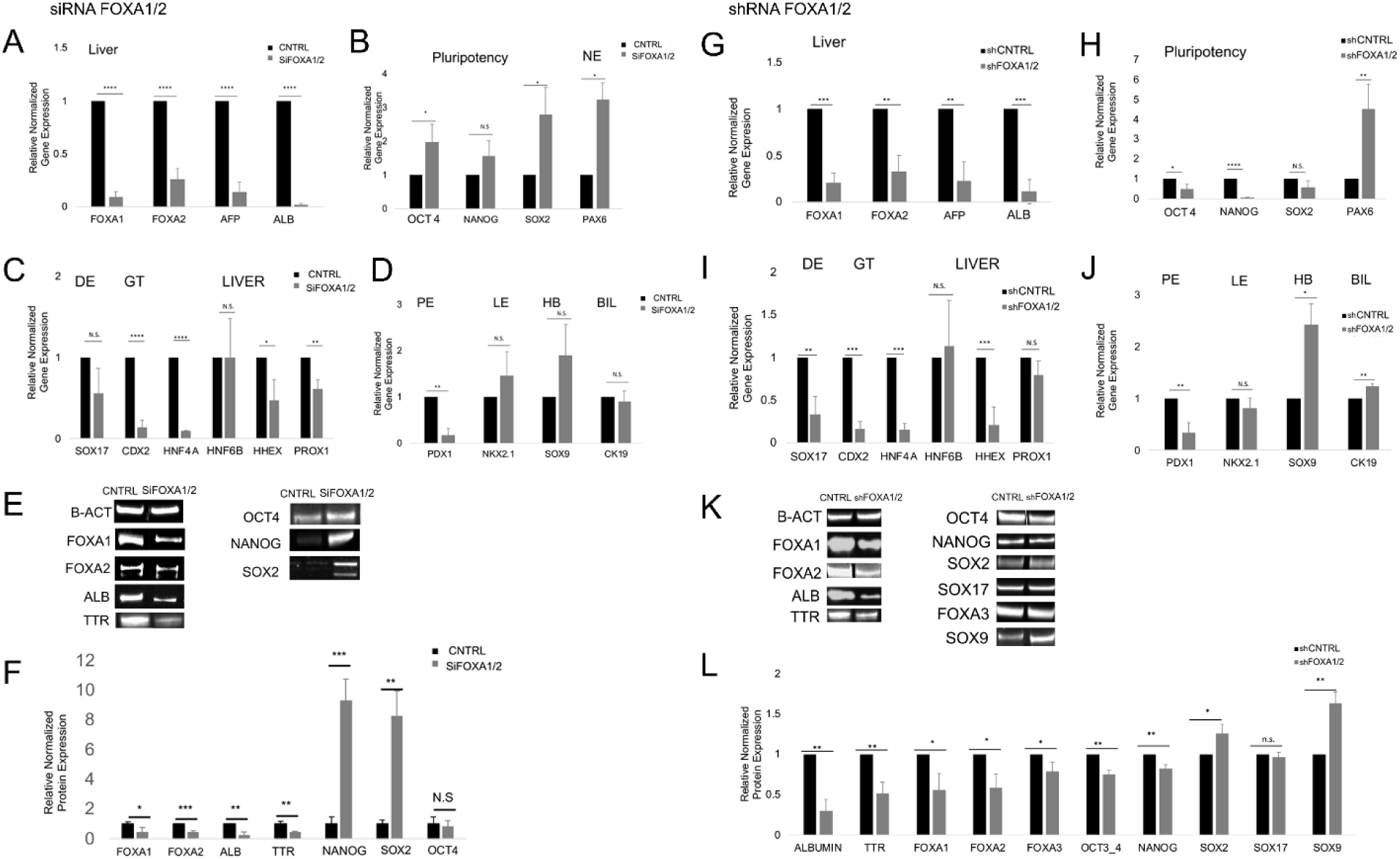
shFOXA1/2 phenotype in spontaneously differentiated human stem cell-derived hepatic progenitors. A) Bar graph of gene expression (qRT-PCR) of Liver genes (FOXA1/2, AFP, ALB) on Day 14, after siRNA scramble and siFOXA1/2 knockdown on Day 11 of culture, for GF (-) condition. Plotted is mean ± SD. Significance defined as *, P < 0.05, **, P <0. 01, ***, P < 0.001. (n = 3). B) Same as L) except pluripotency genes analyzed. C) Same as L) except DE, GT, liver genes tested. D) Same as L) except pancreatic endoderm (PE), lung endoderm (LE), hepatobiliary (HB), biliary (BIL) genes tested. E) Western blot on Day 14 of GF (-) condition, after siRNA FOXA1/2 transduction on day 11 hepatic progenitor cells in the GF (-) condition, in Control (scrambled) and shFOXA1/2 condition. GAPDH is control. F) Quantitation of Western blot on left. Significance defined as *, P < 0.05, **, P <0. 01, ***, P < 0.001. (n = 3). O) Bar graph of gene expression (qRT-PCR) after siFOXA1/2 knockdown on Day 11 assessed on Day 14. Significance defined as *, P < 0.05, **, P <0. 01, ***, P < 0.001. (n = 3). G) Bar graph of gene expression (qRT-PCR) of Liver genes (FOXA1/2, AFP, ALB) on Day 14, after shRNA scramble and siFOXA1/2 knockdown on Day 11 of culture, for GF (-) condition. Plotted is mean ± SD. Significance defined as *, P < 0.05, **, P <0. 01, ***, P < 0.001. (n = 3). H) Same as R) except pluripotency genes analyzed. I) Same as R) except DE, GT, liver genes tested. J) Same as R) except pancreatic endoderm (PE), lung endoderm (LE), hepatobiliary (HB), biliary (BIL) genes tested. K) Western blot on Day 14 of GF (-) condition, after siRNA FOXA1/2 transduction on day 11 hepatic progenitor cells in the GF (-) condition, in Control (scrambled) and shFOXA1/2 condition. GAPDH is control. L) Quantitation of Western blot in K).

Next, we performed lentiviral-mediated shRNA FOXA1/2 knockdown (as opposed to siRNA) during liver differentiation on day 11 and assessed cells on day 14. Interestingly, we once again observed a significant downregulation of liver differentiation genes **(Fig. 7G).** However, in contrast to the siRNA FOXA1/2 phenotype, we observed a downregulation of pluripotency genes OCT4 and NANOG, with SOX2 trending downwards **(Fig. 7H).** We again observed an upregulation of the neuroectodermal marker PAX6 **(Fig. 7H)**. We also observed a significant downregulation of the endoderm marker is SOX17 and the gut tube endoderm CDX2 **(Fig. 7I)**. We again observed downregulation of the liver differentiation markers HNF4A, hHEX, PROX1, but no change in HNF6B **(Fig. 7I)**. Finally, we observed downregulation of the pancreatic endoderm marker PDX1, and concomitant upregulation of the hepatobiliary markers SOX9, the biliary marker CK19, with no significant change in lung endoderm **(Fig. 7J)**. Our data suggests that sustained expression of shFOXA1/2 as well as siFOXA1/2 results in inhibition of liver differentiation. However, in contrast to the siFOXA1/2 effects, shFOXA1/2 results in a downregulation of SOX17 and pluripotency factors, as well as an upregulation of hepatobiliary markers SOX9 and CK19. We provide a schematic that summarizes our findings in shFOXA2 HepG2 cells **(Fig. 8A)** and in both siFOXA1/2 and shFOXA1/2 treated cells during hepatic differentiation **(Fig. 8B-C)**.

**Figure 8.**
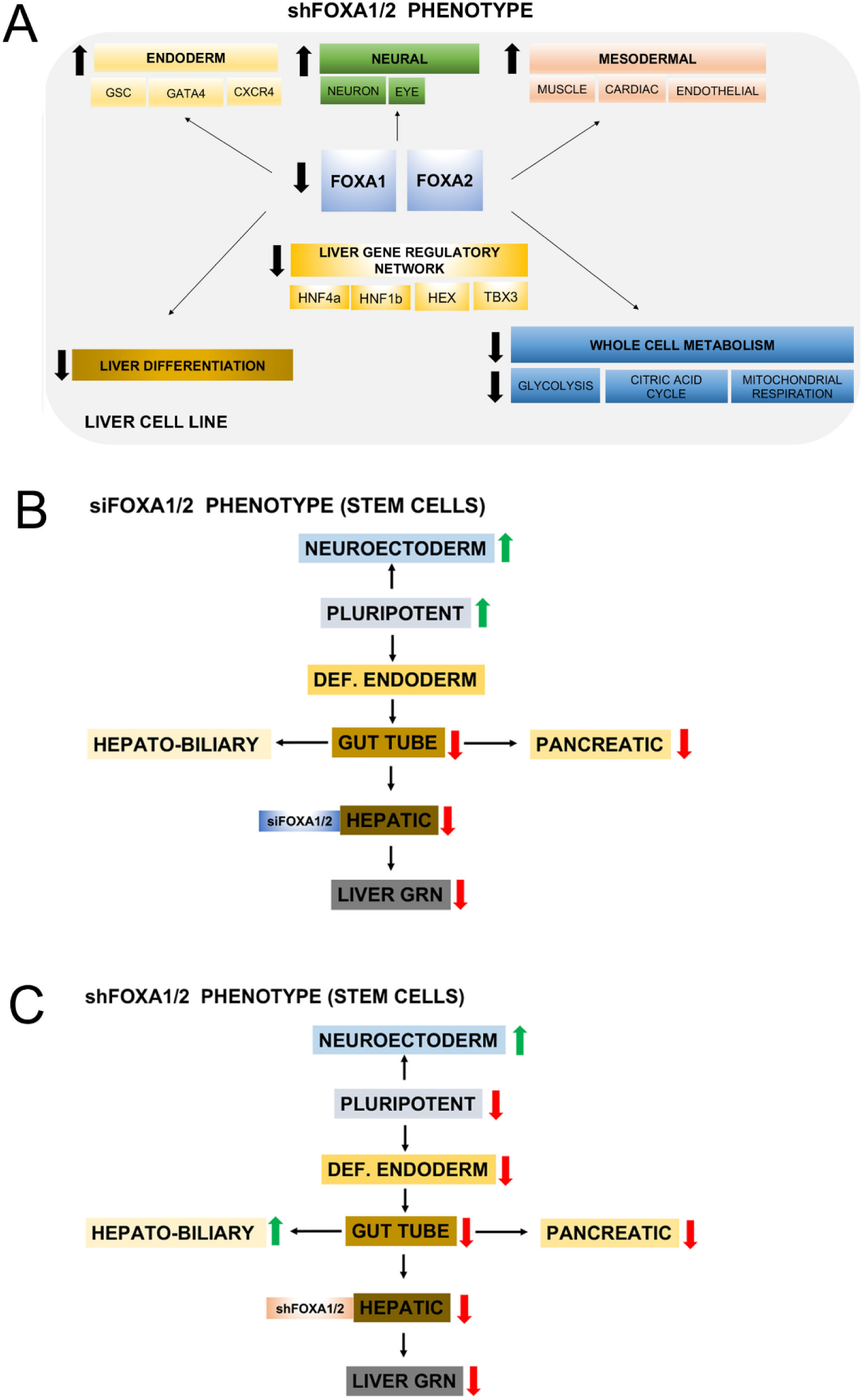
Schematic summarizing FOXA1/2 targeting by RNAi in HepG2 cells and during hepatic differentiation of hPSC. A) A schematic summarizing the shFOXA1/2 phenotype in HepG2 cells based upon qRT-PCR and RNA-seq, and functional analysis (metabolism). Our findings indicate the shFOXA1/2 phenotype results in 1) decrease in both FOXA1/2 (light blue), 2) decreases in the liver gene regulatory network genes (dark yellow), 3) upregulation of genes for endoderm (light yellow), neural (green), and mesodermal genes (pink), 4) downregulation of liver differentiation genes (brown), 5) downregulation of whole cell metabolic pathways (dark blue), including glycolysis, citric acid cycle, and mitochondrial respiration. B) A schematic summarizing gene expression of lineage-specific markers and master transcription factors of specific lineages in day 14 hepatic cells derived from hPSC that have been treated with siFOXA1/2. Upregulation of master factors of Neuroectoderm (PAX6) and pluripotency. C) Same as B except shFOXA1/2 treatment. Findings include an upregulation of neuroectoderm (PAX6) and hepatobiliary markers (SOX 9, CK19).

## DISCUSSION

How developmental GRN TFs control differentiation and metabolism remains a central question in cell, developmental, stem cell, and cancer biology, and is crucial for understanding physiological and pathophysiological processes. FOXA2 is a well-studied member of the hepatic GRN and has a major role in controlling murine liver development and liver metabolism, in part through controlling other GRN members. Studies of the role of FOXA2 in human liver cells are lacking compared to mouse, and the role for FOXA1/2 in human liver differentiation is unclear. We employed siRNA and shRNA to target both FOXA1/2 in both a human liver cell line, and during hPSC-hepatic differentiation. differentiated towards the liver lineage. In a well-studied liver cell line (HepG2, hepatoblastoma cell line), we observed that both siFOXA1/2 and shFOXA1/2 knockdown results in widespread downregulation in liver gene expression. We observed downregulation of FOXA1/2, HNF4A, HNF1B, hHEX, TBX3 in shFOXA1/2 HEPG2 cells, and RNA-seq/bioinformatics analysis demonstrated coordinated upregulation of cell differentiation genes associated with other lineages, including mesoderm-derivatives (cardiac, muscle, endothelial cells) and neuroectoderm. We also identified global and widespread changes in genes associated glycolysis and glucogenesis, the citric acid cycle, and oxidative phosphorylation. Our functional analysis of shFOXA1/2 knockdown cells demonstrated significant changes in both glycolytic and mitochondrial functions, and changes in gene expression of FOXA1/2-dependent glycolytic enzymes. We targeted liver GRN with siFOXA1/2 and shFOXA1/2 during hPSC-hepatic differentiation. The data demonstrated a downregulation of liver differentiation, including a downregulation of all major liver GRN TF, as well as upregulation of neuroectoderm, and pluripotency genes (siRNA) or (shRNA). Our analysis also demonstrates unique lineages that arise during liver de-differentiation, thereby establishing novel regulatory relationships between FOXA1/2 and liver GRN, as well as TF in DE, GT and its derivatives, biliary cells, and pluripotency. Taken together, these data suggest that FOXA1/2 knockdown results in global changes in cell state associated with loss of liver differentiation, gain of differentiation of alternate lineages, and inhibition of both glycolysis, TCA cycle, and oxidative phosphorylation. These findings clearly establish the role of FOXA1/2 in liver cells and should further our knowledge regarding the role of FOXA1/2 and liver GRN in stem cell differentiation, metabolism, liver regeneration, and disease states like cancer and liver fibrosis.

It is important to highlight our findings regarding the initiation of human liver differentiation in our hPSC liver differentiation model. Since we observe a stark downregulation of TF that compose liver GRN, and liver differentiation genes, our data strongly suggests that FOXA1/2 initiates human liver differentiation by activating hepatic GRN in addition to directly activating liver specific genes. Further, we noted an upregulation of pluripotency genes OCT4, SOX2, and NANOG (trending) in the case of the siFOXA1/2 phenotype. Thus, the hPSC express pluripotency genes initially, then downregulate them in DE, and then upregulate them upon siFOXA1/2 silencing in hepatic progenitors. Interestingly, it has been shown that OCT4 and SOX2 knockdown activate endoderm TF ^62^. This suggests a complex relationship between FOXA1/2 and pluripotent markers in early hepatoblasts. Our data suggests that early hepatoblasts are sensitive to FOXA1/2 fluctuation, which could cause transient changes in differentiation status. Some important remaining questions are whether in more mature hepatoblasts, or hepatocytes, whether FOXA1/2 silencing can activate pluripotency factors. Paradoxically, the shFOXA1/2 phenotype resulted in downregulation of OCT4 and NANOG, suggesting sustained silencing causes forward differentiation, further highlights the complexity of GRN relationships. Additionally, SOX9, and CK19 were upregulated, suggesting perhaps biliary differentiation was activated upon de-differentiation. SOX9 has been linked to hepatic, biliary, and pancreatic tissue regeneration, as a marker for undifferentiated precursors in the biliary tract ^63^. Interestingly, FOXA2 and GATA6 have co-occupancy at the SOX9 locus, supporting FOXA2 regulation ^64^. It should be noted that both siFOXA1/2 and shFOXA1/2 phenotypes also demonstrated an upregulation of PAX6, the neuroectodermal master factor which we also observed in the HEPG2 studies. Since we observed upregulation of SOX2, and SOX2 and PAX6 upregulation are associated with neural differentiation ^65^, it is possible that a portion of cells also differentiated along the neuroectodermal lineage. Future studies can determine the differentiation potential of these cells to distinguish between biliary and neural differentiation, and further understand the de-differentiation process.

One of the main findings her was the identification that FOXA1/2 downregulates liver GRN and liver-specific genes in a liver cell line. The siRNA and the shRNA FOXA1/2 knockdown phenotype in the liver cell line (HepG2) has not been previously published. This phenotype demonstrated potent effects on liver GRN, including HNF4A, HNF1B, HNF6, HEX, and TBX3, which is a non-obvious result, since FOXA1 and FOXA2 knockout has minimal effects on late-stage fetal hepatocytes *in vivo* ^22^, even though HepG2 cells resemble de-differentiated hepatocytes. Although the HepG2 cells are tumor-derived (Hepatoblastoma), it is possible that GRN and their downstream functions are preserved. Therefore, our findings represent a form of a hepatic differentiation collapse, as has been reported recently (FOXA1/2/3 (-/-/-) phenotype) ^16^. It is important to note that RNAi of FOXA1/2 has been employed in human liver cell lines for analysis of gene regulation of AFP ^40^ and lipoprotein lipase ^66^. These studies both employed siRNA targeting FOXA2 in human cell lines, but not both FOXA1 and FOXA2. Kanaki *et al*. demonstrated effects of FOXA2 downregulation on downregulation of ALB and transferrin, as well HNF4A and HNF1B in HepG2 cells, but reports a decrease in HNF6, which we did not observe. Their investigation demonstrated that they also inadvertently targeted the FOXA1 gene. However, these studies did not examine hepatic GRN, lineage-specific markers, both siRNA and shRNA phenotypes, and reversibility. Interestingly, FOXA2 knockout in adult livers (by E18.5), resulted in no functional consequence ^67^, probably due to compensation by FOXA1, while FOXA1/2 knockout in embryonic liver at E16.5 exhibited phenotypic consequences in the biliary tract, and did not demonstrate any changes in hepatocyte morphology, ultrastructure, or function, other than a 10-fold increase in AFP expression ^68^. Overall, our data in the FOXA1/2 depleted HepG2 cells is partially consistent with the Riezel *et al.* study, in that we observe downregulation of cytochrome P450 genes and apolipoproteins (**Table 3, Supp. File 4**), as well as ALB, and is consistent with a differentiation collapse in HepG2 cells. Interestingly, in our shFOXA1/2 knockdown cells, we observed upregulation of GATA4, which is not only an endoderm and mesoderm marker, but also displays pioneer factor activity at ALB (liver) genes ^19^.

Although FOXA3 is believed to compensate for the loss of FOXA1/2, our studies indicate that the relationship between FOXA1/2 and FOXA3 expression is complex. For example, our analysis of the siFOXA1/2 phenotype indicates that FOXA3 gene expression was downregulated **(Fig. 1I),** whereas in the the shFOXA1/2 phenotype, FOXA3 was upregulated **(Fig. 2D)**, even though both showed a strong de-differentiation phenotype. In light of this complex relationship, we are currently performing triple knockdown experiments and will publish this work in a future manuscript. The Riezel et al. study showed that histone modifications (H3K27) which predict gene activity are affected by the absence of FOXA factors, suggesting FOXA1/2 factors are required to maintain epigenetic states in adult cells, but it is unclear how epigenetic mechanisms play a role in our study. Future investigation employing CRISPR-based gene knockout strategies are needed to fully understand the FOXA1/2/3 knockdown phenotype.

An interesting aspect of the shFOXA1/2 phenotype is the upregulation of genes for endodermal (endoderm, gut), mesodermal (cardiac, endothelial, muscle) and neuroectodermal (neural, eye, skin) morphogenesis **(Fig. 4D)**. To our knowledge, this phenotype in a de-differentiated hepatic carcinoma (hepatoblastoma or hepatocellular carcinoma) has not been identified, and may represent de-differentiated hepatoblastoma or embryonal carcinoma cell type. One possibility is that diminished FOXA1/2 in a de-differentiated HepG2 cell results in decreased downstream DNA binding, which results in activation of earlier lineages (endoderm) and alternate lineages (gut, mesoderm and neuroectoderm). This data is consistent with models of GRN and how cell lineages are formed as a function of regulatory TFs ^69^. In this model, the establishment of endoderm and eventually liver would result in repression of master TFs associated with mesoderm (cardiac, endothelial cells, muscle) and neuroectoderm (neural, eye, skin). Interestingly, studies suggest that FOXA2 positively regulates cardiac differentiation/morphogenesis at its earliest stages ^70–72^. Similarly, FOXA2 regulates neuroectoderm and neural differentiation ^73–76^. However, we found that the loss of FOXA2 appears to increase cardiac and neural gene expression **(Fig. 4D)**, which suggests negative regulation, not positive regulation. Therefore, in HepG2, the loss of FOXA1/2 and the hepatic GRN TF leads to a direct or indirect loss of repression of cardiac or neural genes, together with an activation of endoderm genes. Interestingly, genetic studies of conditional knockout of Foxa1/2 within pancreatic islet cells demonstrates not only loss of the islet phenotype, but also activation of neural genes, including Tacr3 (dopaminergic activity), and other neural genes ^52^. Thus, in both hepatic and pancreatic cells, FOXA1/2 negatively regulates neuroectoderm/neural differentiation. Another possibility, is that hepatic de-differentiation leads to activation of pancreas differentiation at the gene regulatory level, leading to enhanced neural gene expression. We speculate that phenotypes of de-differentiation and alternate lineage activation can be explained by studies of hepatic GRN. FOXA2 is known to initiate FOXA1, HNF1B, and HNF4A expression, and thereby control differentiation and metabolism in visceral endoderm derived from mouse embryonic stem cells ^20^. In human stem cell models, other putative activators of hepatic GRN include GATA6 ^77^ and HNF4α ^78–79^, which could help explain the phenotypes that we observe. The increase in CDX2 (gut differentiation) could be explained by the loss of FOXA2 repression, as reported in murine studies ^80^. Based on above, the shFOXA1/2 phenotype in HepG2 cells shares similarity with hPSC-derived hepatocyte-like cells (Raju, Chau et al. 2018). Interestingly, this meta-analysis study shows that: 1) the resulting hPSC-hepatocyte-like cells in the field are fetal-like and not fully mature, 2) they express a set of undesired genes, including cardiac mesoderm and neural/eye development, 3) they are not fully functional metabolically. We speculate that FOXA1/2 dysregulation may affect hPSC differentiation.

Our major findings in the shFOXA1/2 knockout are that glycolysis, TCA cycle, and mitochondrial metabolism are all highly downregulated. A major question is how do these cells generate energy? Liver cell lines like HepG2 cells have been widely used for human liver metabolism and xenobiotic metabolism ^81–86^ . In previous studies of HepG2 metabolism, a “no glucose” condition exhibits very low glycolysis, which may also model blocked glycolysis ^81^. In this model of HepG2 glycolysis inhibition, energy generation was generated by glycolysis (NADH, ATP), and glutaminolysis (NADPH and NADH via TCA cycle), pyruvate oxidation (NADH), oxidative phosphorylation (ATP), and conversion of proline to glutamate (NADPH) in the urea synthesis pathway ^81^. Our system differs, because pyruvate dehydrogenase complex subunits are downregulated, which would reduce pyruvate oxidation, in addition to the deficit we observe in mitochondrial metabolism. Interestingly, chemical inhibition of glycolysis and oxidative phosphorylation via 3BrPA (3-bromopyruvate) ^87^ mimics the phenotype we observe in shFOXA1/2 cells. These chemical inhibition studies demonstrate key insights to carbon sources for energy production. This is an effective strategy to treat hepatocellular carcinoma (HCC), while sparing normal tissue in a rabbit hepatoma model ^88^. Other studies have employed metabolic targeting in cell death ^89^ and sensitivity to chemotherapeutic treatment ^87^. While 3BrPA achieves inhibition of glycolysis and mitochondrial metabolism, to our knowledge, this is the first report of mitochondrial dysfunction being attributed to FOXA1/2 knockdown. We hypothesize this is due to the coordinate downregulation of PPARGC1a, a transcriptional coactivator of mitochondrial metabolism ^55–56^. PPARGC1a exerts its effect on mitochondrial metabolism, including binding to retinoid receptors (RXR), farnesyl X receptors (FXR), pregnane receptors (PXR), liver receptor (LXR), HNF4a, estrogen related receptor (ERR), and others ^55^. Thus, glycolytic and mitochondrial defects in shFOXA1/2 cells can be used for designing metabolic treatment for HCC. Further metabolic studies can help identify sources of cellular energy and metabolic fluxes in shFOXA1/2 cells. Regarding nitrogen metabolism, HepG2 cells follow alternate urea/arginase pathways which could employ external sources of glutamine for glutaminolysis, and the arginase II pathway via putrescine metabolism for alpha-keto glutarate, and succinate generation, which would enable anabolic reactions and cellular energy generation ^81, 90^ . Since HepG2 cells are tumor-derived and therefore have higher rates of glycolysis and glutaminolysis, and lower rates of oxidative phosphorylation than primary hepatocytes, suggesting alternate energy generation to oxidative phosphorylation ^81^. This is consistent with metabolic reprogramming, which includes altered bioenergetics, biosynthesis, and redox balance ^91^. Another potential mechanism is that oncometabolites that can feedback to alter gene expression and cell fate ^91^.

Thus, shFOXA1/2 knockdown combined chemical or genetic targeting of tumor metabolism could be a method for elucidating tumor metabolism. In fact, glycolysis inhibitors are a real pharmaceutical strategy for treating HCC ^92^ and combining glycolysis and mitochondrial approaches is considered a synergistic approach for treating HCC ^93^. Interestingly, glioblastomas have been shown to utilize acetate rather than glutamate as a carbon source ^57^, through upregulation of acetyl-CoA synthetase enzyme 2 (ACSS2). Here, we observed ACSS2 expression in the shFOXA1/2 knockdown cells, which suggests may employ acetate as carbon source. Further characterization of the FOXA1/2 in primary cells and tumors should elucidate these important questions.

Our findings in shFOXA1/2 regarding both differentiation and metabolism effects could be applicable to a wide range of tumors derived from human tumors. Although FOXA1/2 mutations have not been directly linked to hepatocellular carcinoma (HCC) ^94^, FOXA dysregulation controls either tumor suppressors or oncogenes during HCC tumorigenesis ^32^ and overall, FOXA genes effect many hallmarks of tumorigenesis ^95^. In human lung cancer, FOXA2 is suppressed, whereas FOXA1 and FOXA3 are increased ^96^, and FOXA3 is a potential biomarker. Further, FOXA1/2 is involved in lineage switching from lung to gastric identities in mucinous lung cancer with temporal effects over cell fate ^97^. In endocrine-resistant breast cancer, FOXA1 over-expression reprograms endocrine-resistant breast cancer, resulting HIF-2A mediated metastasis ^98^. FOXA1 has been shown to participate in epigenetic mechanisms in liver, breast, and prostate in cell lines ^33^. FOXA factors regulate epithelial mesenchymal transition (EMT) in colon cancer through pioneer factor function ^99^. In prostate cancer, FOXA1 mutations effect tumorigenesis, with a prevalence of ∼35%, including separate classes of activating mutations that transactivate androgen receptor, enhance metastasis, and drive oncogenes ^100^. Finally, FOXA1/2/3 have all been shown to be involved in development and progression of bladder cancer ^101^. We speculate that when FOXA expression may cause tumorigenesis, it may be accompanied by the activation of genes for alternate lineages, and metabolic programming which may hinder therapeutic targeting.

There are limitations to our study worth mentioning. We employed RNAi because of its flexibility and ease of use. However, we are not obtaining the complete FOXA1/2 (-/-) phenotype, which can be implemented with CRISPR knockout of FOXA1/2 in liver cell lines, hPSC-derived hepatocytes, hepatocytes, and primary liver cancer cells. Nonetheless, downregulation of FOXA1/2 may be more valuable for modeling disease phenotypes like nonalcoholic liver disease ^50^ and fibrosis ^102^. Another limitation is regarding our approach for knockdown. We employed antibiotic-based selection to generate our stable cell lines of interest, both with lentivirus (shRNA) and nonvirally (conditional cell line). While it is possible that antibiotic selection resulted in some changes in cell change phenotype, we did not observe any measurable changes. Nonetheless, future studies will employ reporter-based strategies can be performed with ease and less perturbation of cell state. Although we focused on a liver cell line here, and it has been widely studied, the cells exhibit genetic alterations and adaptations to cell culture. Therefore, future studies will focus on phenotypes in human hepatocytes and human primary liver cancer, which may improve the scientific relevance of our work. Another aspect of our analysis is that we employed a polyclonal population with respect to lentiviral transduction and insertion of the shFOXA1/2 genes. This protects against promoter shut down or against insertion effects, because in a polyclonal population these effects would be minimized. On the other hand, single cell clones may provide more detailed information and could be obtained through single-cell cloning procedures. Although we don’t expect multiple subpopulations in our shFOXA1/2 cells, single cell cloning as well as single cell-RNAseq may provide more answers to whether multiple subpopulations exist, which would indicate a more complex response to shFOXA1/2 knockdown. Finally, the relationship between the shFOXA1/2 knockdown phenotype and expression of HNF1A and HNF1B needs to be further delineated, in terms of its role in liver differentiation and metabolism. HNF1B is normally detected during early gut development, and both HNF1A/B are expressed in the developing liver ^1^, and HNF1A is expressed in the adult liver. We detected decreases is HNF1B in the shFOX1/2 cells, but we did not detect changes in HNF1A expression. It is unclear whether HNF1B was compensating for HNF1A functions, as has been reported ^103^. HNF1A also has widespread metabolic effects in the liver ^1^ and its role in the shFOXA1/2 knockdown cells needs to be determined. Here, we have provided evidence that hepatic GRN networks can be controlled by FOXA1/2 in human stem cells, at the level of endoderm induction and hepatic endoderm differentiation. Moreover, we find that FOXA1/2 phenotype in stable liver cells (HepG2), a key model for liver GRN studies, has potent, genome-wide effects in blocking liver differentiation and seemingly reversible effects on cells. Further we find that siFOXA1/2 regulates pluripotency factors, biliary markers, and neurectoderm during hPSC liver differentiation. It remains to be seen how our understanding of GRN can enable the improved differentiation and maturity of hepatic progenitors, and of alternate cell fates like intestine, pancreas, thyroid, and lung. Further understanding of GRN, like compensatory mechanisms, concentration effects, and reversibility will likely alleviate challenges in engineering transcriptionally complex hepatocytes from hPSC and help coordinate appropriate epigenetic and genetic changes that occur during differentiation.

## LIST OF ABBREVIATIONS

AFP: alpha-fetoprotein Dox doxycycline
ECAR: extracellular acidification rate EGF epidermal growth factor
EMT: epithelial mesenchymal transition FBS fetal bovine serum
FDR: false discovery rate FOXA Forkhead box A
GDR: gentle cell dissociation reagent GO gene ontology
GRN: gene regulatory networks GSEA gene set enrichment analysis HCC hepatocellular carcinoma HepG2 human hepatoma
hESC: human embryonic stem cell HGF hepatocyte growth factor
hPSC: human pluripotent stem cells iPSC induced pluripotent stem cell MTG monothioglycerol
NES: normalized enrichment score OCR oxygen consumption rate P/S penicillin-streptomycin
PCA: principal component analysis PCR polymerase chain reaction RNA-seq RNA-sequencing ROCK rho-associated kinase
RT: reverse transcription siRNA short-interfering RNA TFs transcriptional factors

## Competing Interests

The authors hereby state no competing interest involved with the ideation, writing, or revision of this manuscript

## ACKNOWLEDGEMENTS

The authors acknowledge the UB Center for Excellence in Bioinformatics, and Roswell Park Lentiviral core facility. NP was supported by the UB CBE startup funds, and by the UB Center for Cell, Gene and Tissue Engineering (CGTE). NP and MM were supported by the New York State Stem Cell Science C024316, and the Stem cells in regenerative medicine (ScIRM) center. OO was supported by the Western NY Prosperity Fellowship. The authors acknowledge that the shFOXA2 vectors were a kind gift obtained from Professor Jianguo, Yunneng Tang, and Weiwei Shi, Institute Pasteur of Shanghai, Chinese Academy of Sciences, Shanghai, China. We acknowledge that our article has been accepted as a preprint at https://www.biorxiv.org/content/10.1101/2020.06.01.128108v2.

## AUTHOR CONTRIBUTIONS

IY: Obtained data, analyzed data, built figures, wrote and approved manuscript

MM: Obtained data, analyzed data, built figures, edited and approved manuscript

DG: Obtained data, analyzed data, built figures, approved manuscript

AC: Obtained data, analyzed data, built figures, approved manuscript

MM: Obtained data, analyzed data, built figures, edited and approved manuscript

OO: Obtained data, analyzed data, built figures, approved manuscript

TG: Obtained data, analyzed data, built figures, approved manuscript

TM: Obtained data, analyzed data, built figures, approved manuscript

SM: Obtained data, analyzed data, built figures, approved manuscript

XL: Obtained data, analyzed data, built figures, approved manuscript

ST: Obtained data, analyzed data, built figures, approved manuscript

AS: Obtained data, analyzed data, built figures, approved manuscript

RT: Obtained data, analyzed data, built figures, approved manuscript

PC: Obtained data, analyzed data, built figures, approved manuscript

RP: Obtained data, analyzed data, built figures, approved manuscript

MY: Conceptualized, acquired funding, investigated, edited manuscript, and approved manuscript.

SK: Conceptualized, acquired funding, investigated, supervised, wrote, edited manuscript, and approved manuscript.

NP: Conceptualized, acquired funding, investigated, supervised, wrote, edited manuscript, and approved manuscript.

## DATA AVAILABILITY STATMENT

The transcriptomic data that support the findings in this study are openly available on the gene expression Ominibus. All other data are included within the manuscript.

### Compliance with Ethical Statements

Conflict of Interest: The authors have no conflicts of interest regarding this manuscript.

Funding: This study was undertaken with startup funds, support of the UB Stem cells in regenerative medicine (ScIRM) center (NYSTEM C024316), and the UB Center for Cell, Gene and Tissue Engineering (CGTE).

Ethical approval: This article does not contain any studies with human participants or animals performed by any of the authors.

## Notes

### Competing Interest Statement

We have applied for intellectual property.

### Summary of Updates

New data and updates to the text

