## Supplementary figures and images for "FOXA1/2 depletion drives global reprogramming of differentiation state and metabolism in a human liver cell line and inhibits differentiation of human stem cell-derived hepatic progenitor cells"

### Supplemental Figure 1

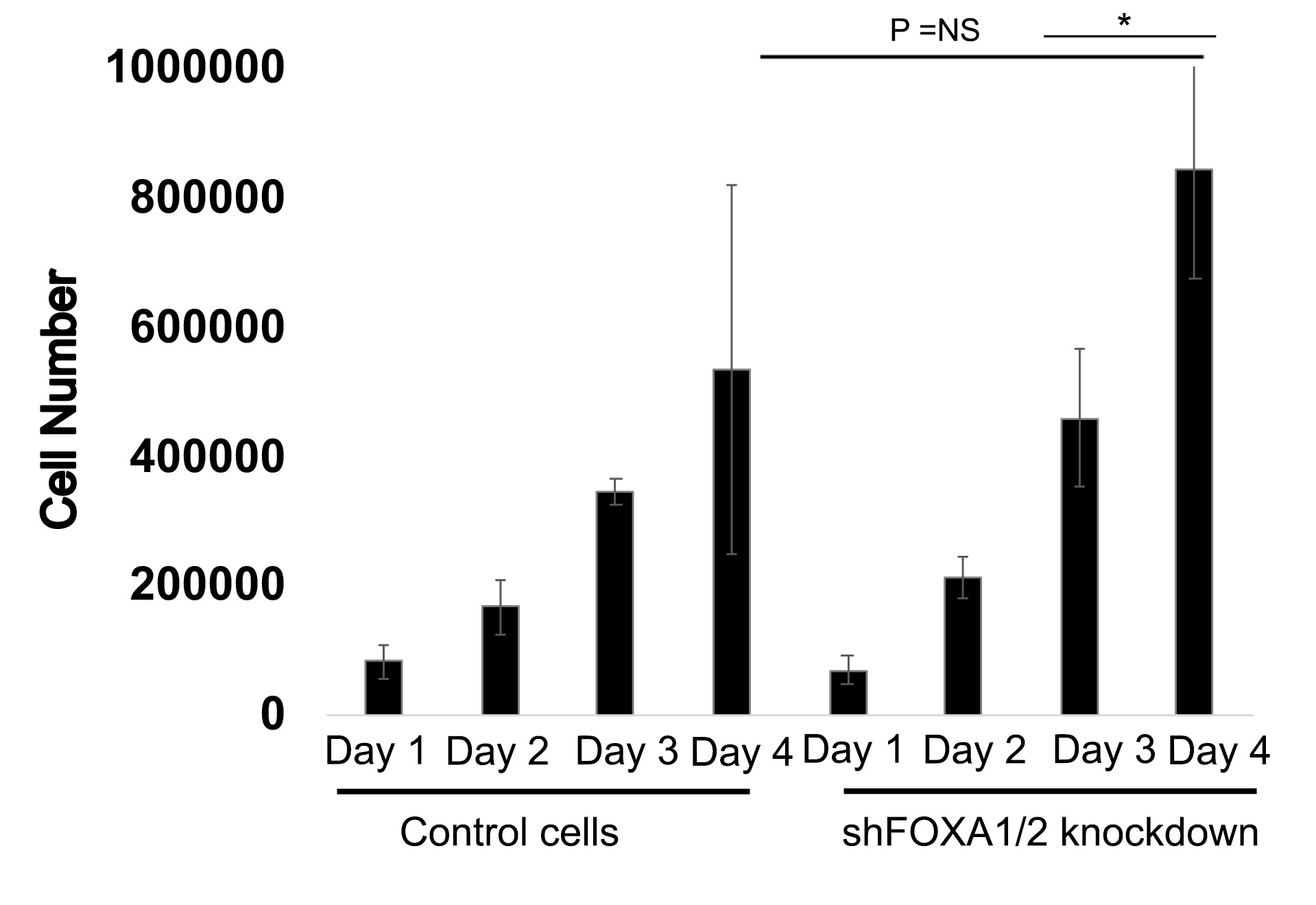
